# Electrically Enhanced Intraperitoneal Drug Delivery Resolved by Fluorescence Lifetime Imaging Microscopy

**DOI:** 10.64898/2026.10.29.762719

**Authors:** Nidda Saeed, Irina A. Okkelman, Sergey M. Borisov, Dauwe Bulckaen, Ruslan I. Dmitriev, Wim Ceelen

**Author notes:** Shared senior authorship (Ruslan I. Dmitriev, Wim Ceelen).

## Abstract

Intraperitoneal delivery of nanoparticles (NPs) holds promise in the treatment of peritoneal metastases (PM), and the use of electric fields may enhance their tissue penetration. Evaluating NP delivery requires evidence of intracellular payload release, which is challenging in autofluorescent tissues. Here, we combined electromotive drug administration (EMDA) with cationic RL-100 NPs co-loaded with doxorubicin (DOX) and a near-infrared aza-BODIPY tracer, reporting carrier-associated distribution, while phasor fluorescence lifetime imaging microscopy (FLIM) characterized the molecular environment of DOX. In colorectal carcinoma, NP-delivered DOX progressed from cytoplasmic puncta to nuclear accumulation, while carrier-associated fluorescence remained cytoplasmic, supporting intracellular release. EMDA enhanced NP penetration in ex vivo human peritoneum and healthy rat peritoneum. In a syngeneic rat model of colorectal PM, both free and NP-delivered DOX remained detectable to approximately 1200 µm after EMDA, compared with < 200 µm after passive delivery. NP-delivered DOX remained detectable 24 h after EMDA. Whole-animal and ex vivo imaging showed the highest carrier-associated signals in tumor nodules and tumor-bearing omentum. These findings support electrically enhanced intraperitoneal delivery and a dual-reporter approach to evaluate carrier distribution, drug penetration, and intracellular release, with potential applications in other autofluorescent tissues. Together, these capabilities provide a preclinical basis for investigating NP-assisted delivery.

## 1. Introduction

Peritoneal metastasis (PM) represents a challenging manifestation of colorectal cancer, being reported in approximately 4–5% of patients at diagnosis. Following curative resection, reported rates of metachronous PM range from 4–12% in colon cancer and 2–19% in rectal cancer. Median survival remains near 23 months despite modern systemic therapy, reflecting the limited access of systemically delivered drugs to peritoneal deposits.^1^ Intraperitoneal (IP) administration places cytotoxic agents in direct contact with tumor deposits, but penetration from the peritoneal surface into tumor tissue remains limited, leaving deeper cell populations underexposed.^2,3^

Elevated interstitial fluid pressure (IFP), outward interstitial flow near the tumor boundary, and a dense extracellular matrix restrict transport into tumor tissue after IP drug delivery.^2,4,5^ In a human clinical series of primary and metastatic breast and colorectal carcinomas, IFP averaged 20 ± 13 mmHg, compared with 2 ± 4 mmHg in normal tissue (mean ± SD).^6^ In tissue-isolated rat tumors, pressure increased from the surface toward the interior, reaching a plateau at depths of 0.2–1.1 mm, the same range over which IP-delivered drug must travel to reach viable tumor.^7^ Nanoparticle (NP) carriers represent a promising IP drug delivery vehicle, prolonging peritoneal residence, reducing systemic exposure, and accommodating various therapeutic and imaging payloads. Thus, formulations ranging from lipid-coated mesoporous silica to enzyme-responsive prodrugs and taxane-loaded systems have shown promise in preclinical peritoneal metastasis models.^8–10^ In mice bearing peritoneal pancreatic carcinoma xenografts, paclitaxel-loaded mesoporous silica NPs increased peritoneal cavity residence time 3.2-fold and peritoneal tumor accumulation 6.5-fold relative to free paclitaxel, while reducing systemic exposure to approximately one-third.^11^ However, prolonged residence within the peritoneal cavity does not necessarily translate into deeper penetration into tumor tissue.

Physical delivery strategies can address this limitation by applying an external driving force.^12^ Electromotive drug administration (EMDA) uses a direct current (DC) electric field to enhance transport of charged species through electromigration and electroosmosis, principles established in transdermal iontophoresis and subsequently applied to other tissues and organs.^13,14^ In oncology, EMDA has demonstrated clinical efficacy in intravesical chemotherapy^15^ and has been investigated experimentally in pancreatic and other solid tumors.^16–18^ Related electrostatic approaches, including charged aerosol delivery, have also improved IP NP deposition.^19,20^ Direct-current electromotive transport had not previously been applied to IP drug delivery. We established this approach in a syngeneic rat model of colorectal PM, demonstrating enhanced penetration of positively charged NPs into healthy peritoneum and tumor nodules and defining the optimal electrical parameters, exposure time, and carrier solution: a maximum tolerated current of 1.6 mA applied for 30 min in 50 mL normal saline.^21^ That work used a model fluorophore that reported only carrier location, so neither drug delivery nor payload release could be assessed.

The challenge of successful detection of drug release requires distinguishing three co-existing signals within a single specimen: intact carrier, released drug, and endogenous tissue autofluorescence. Peritoneal tissue exhibits intense broadband autofluorescence from NAD(P)H, flavins, collagen, elastin, and porphyrins, with combined emission spanning approximately 400–700 nm and overlapping with the 491–648 nm detection window used here for visible-range labels.^22,23^ Wavelength-dependent absorption and scattering can further attenuate excitation and emitted light in intact tissue, complicating intensity-based quantification.^24^

These factors interfere with conventional fluorescence imaging, which traditionally separates fluorescent signals by spectral properties and quantifies them by signal intensity. In addition, while doxorubicin (DOX) remains the most widely investigated payload for targeted delivery,^25^ encapsulated and released DOX show identical excitation and emission spectra, differing only in quantum yield.^26^ Spectral measurements alone therefore cannot readily distinguish carrier-held drug from drug that has reached its molecular target, and intensity-based studies have consequently inferred release rather than demonstrated it.^27^

Fluorescence lifetime imaging microscopy (FLIM) addresses these issues by separating fluorescent species based on excited-state decay time and enabling discrimination between fluorophores with overlapping spectra.^28^ Unlike fluorescence intensity, lifetime is not directly proportional to fluorophore concentration or excitation power, although it can be sensitive to molecular interactions and the local (micro)environment.^29,30^ The phasor approach is a popular FLIM analysis method, representing fluorescence decay in each pixel in a two-dimensional coordinate space without fitting multiexponential decay models, thereby distinguishing fluorescent species with different lifetime signatures.^31^ When applied to nanomedicine, FLIM has resolved DOX release from polymeric, mesoporous silica, liposomal, and gold nanocarriers, and phasor analysis has tracked pH-triggered release in living cells.^27,31–35^ Pairing lifetime discrimination with a near-infrared label addresses both obstacles at once: emission beyond 650 nm lies outside the endogenous emission envelope, while lifetime separates carrier-held from released drug within a single spectral channel.^36,37^ Within available dye chemistries, aza-boron-dipyrromethenes (aza-BODIPYs) combine high quantum yield, photostability, and tunable near-infrared emission and have already been successfully incorporated into polymeric NPs for cell imaging, in vivo molecular imaging, nanoplastics research, and imaging-guided therapy.^38–45^

Our previous EMDA study established enhanced penetration of fluorescent NPs into peritoneal tissue and tumor nodules but did not assess a therapeutic payload.^21^ Here, we combined near-infrared lifetime imaging with EMDA to measure carrier penetration and intracellular drug release within the same model system. Polymeric NPs were prepared by nanoprecipitation^46,47^ and co-loaded with a near-infrared aza-BODIPY tracer^45^ and DOX to provide two reporters with separate spectra and lifetimes. Spectrally distinct carrier and payload signals allowed their distributions to be assessed separately, while the environment-sensitive fluorescence lifetime of DOX provided complementary information on its intracellular fate. Carrier penetration was evaluated in ex vivo human and healthy rat peritoneal tissue, while intracellular DOX release was confirmed in colorectal carcinoma cells. A syngeneic rat model of colorectal PM was used to assess tumor penetration, complemented by whole-animal and ex vivo near-infrared imaging of carrier distribution. EMDA enhanced carrier penetration in peritoneal tissue and extended detectable DOX penetration into tumor nodules, where both carrier and drug signals were observed. We quantified tumor depth profiles from DOX fluorescence intensity, while FLIM and intracellular localization provided complementary evidence consistent with payload release. Together, these experiments extend the evaluation of electrically enhanced NP delivery beyond carrier accumulation to include drug penetration and intracellular payload behavior.

## 2. Results and Discussion

### 2.1. Design, Photophysical Properties, and Environmental Response of DOX/aza-BODIPY Co-Loaded RL-100 Nanoparticles

RL-100 NPs were co-loaded with DOX and diBuO-aza-BODIPY to generate a cationic dual-reporter formulation. diBuO-aza-BODIPY served as the carrier label because its near-infrared emission reduces overlap with tissue autofluorescence and is spectrally separated from the visible fluorescence of DOX. DOX is a small amphiphilic anthracycline with a daunosamine amino group (pKa ≈ 8.3) which is predominantly protonated at physiological pH, resulting in a net positive charge. Nanoprecipitation entrapped the comparatively hydrophilic drug and the hydrophobic tracer within the same polymer matrix during solvent displacement (Figure 1a). Previously, transmission electron microscopy of aza-BODIPY-labeled particles without DOX showed a mean particle size of 58.4 ± 3.5 nm (mean ± SEM; n=150 NPs; Figure S1).^45^ Hydrodynamic size and zeta potential could not be measured directly since diBuO-aza-BODIPY strongly absorbs the 633 nm laser emission line of the instrument. However, previously characterized RL-100 formulations prepared by the same procedure of nanoprecipitation, either undoped or doped with dyes that do not absorb in the red region, reported a diameter of 35–70 nm and a zeta potential of +32 to +58 mV,^46,47^ consistent with the presence of positively charged quaternary ammonium groups.^44,45^ This cationic character motivated the choice of RL-100 for electromotive delivery, shown to enhance the penetration of positively charged NPs into peritoneal and tumor tissue.^21,48^

**Figure 1.**
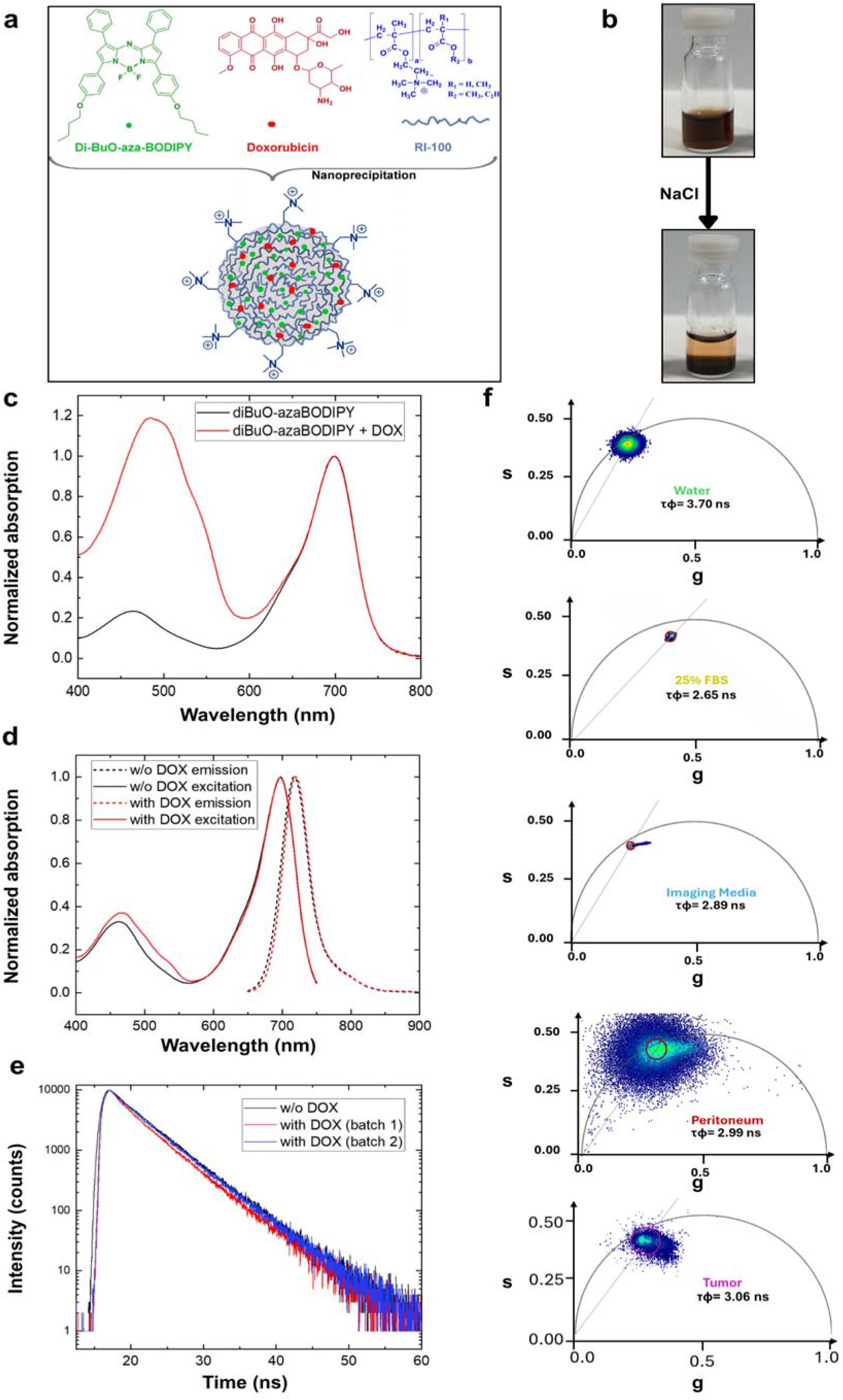
Design, optical characterization, and environment-dependent fluorescence lifetime of DOX/diBuO-aza-BODIPY co-loaded RL-100 NPs. a) Preparation of cationic RL-100 NPs co-loaded with DOX (red) and diBuO-aza-BODIPY (green) by nanoprecipitation. The NP representation is schematic and not drawn to scale. b) Salt-induced precipitation of DOX/diBuO-aza-BODIPY co-loaded NPs before (top) and after (bottom) addition of saturated NaCl (1:5 v/v). c) Absorption spectra of NPs containing diBuO-aza-BODIPY alone (black) or with DOX (red), normalized to the aza-BODIPY absorption band. Dye and DOX inputs were 1 and 10 wt%, respectively, relative to polymer mass. d) Normalized excitation (solid; λem = 770 nm) and emission (dashed; λexc = 635 nm) spectra of NPs without DOX (black) and with DOX (red). e) Fluorescence decays of diBuO-aza-BODIPY in NPs without DOX (black) and in two independent DOX-containing batches (red, blue), recorded at λexc = 635 nm and λem = 720 nm. Fitted lifetimes were 4.4 ns without DOX and 4.2 and 4.4 ns with DOX. f) Phasor plots of diBuO-aza-BODIPY-labeled NPs without DOX in deionized water, 25% fetal bovine serum, imaging medium, fresh human peritoneum, and rat tumor tissue. Labels indicate τφ derived from each phasor position. Phasor data were acquired at 37 °C with an 80 MHz excitation repetition rate.

Co-loading of DOX was confirmed spectrophotometrically. Because RL-100 NPs scatter light weakly, absorption spectra could be recorded essentially free of scattering background (Figure 1c). NPs doped with 1 wt% diBuO-aza-BODIPY alone showed the expected absorbance bands at 464 and 699 nm, whereas NPs additionally loaded with 10 wt% DOX displayed a strong additional band at approximately 480 nm. Based on the molar absorption coefficients of DOX (ε ≈ 13,500 L mol^−1^ cm^−1^) ^26^ and of structurally analogous alkoxy-substituted aza-BODIPY dyes (ε ≈ 80,000 L mol^−1^ cm^−1^),^42^ together with their molecular weights (543 Da for DOX free base and 640 Da for the dye), the DOX-to-dye mass ratio in the dispersion was estimated to be close to the 10:1 ratio used during preparation. These results supported the presence of both components in approximately the expected proportions, although absorption measurements alone did not establish their encapsulation efficiency.

Payload retention was assessed by salt-induced precipitation (Figure 1b). Addition of saturated sodium chloride (1:5 v/v) to a concentrated dispersion (∼5 mg mL^−1^) completely precipitated the particles and left any released payload in the supernatant (Figure S2). Immediately after preparation, the supernatant above the precipitated DOX-containing NPs appeared colorless, indicating complete incorporation of DOX. After prolonged storage in water at 4 °C (∼18 months), the supernatant showed only a faint brown coloration, consistent with minimal DOX leakage, while the hydrophobic diBuO-aza-BODIPY was fully retained. The RL-100 matrix therefore encapsulated the amphiphilic, comparatively hydrophilic DOX efficiently and showed stability during long-term aqueous storage.

DOX co-loading had little effect on the photophysical properties of the tracer, a prerequisite for its use as a lifetime reference. Normalized emission and excitation spectra of the co-loaded NPs closely matched those of the singly doped NPs, with an emission maximum at 718 nm and excitation maxima at 463 and 697 nm, in agreement with the absorption spectra (Figure 1d). Fluorescence decays were described by a mono-exponential function in both cases and were only marginally affected by DOX (τ = 4.4 ns without DOX; 4.2 and 4.4 ns for two independent DOX-loaded batches; Figure 1e), excluding significant Förster-type quenching of the tracer by the co-encapsulated drug. The near-infrared channel thus provided an independent optical reference for the carrier, allowing its distribution to be tracked alongside that of DOX. This distinguishes the dual-label configuration from single-label DOX carriers, in which encapsulated and released DOX contribute to the same fluorescence channel.^27,34^

The fluorescence lifetime of the NP-associated tracer showed sensitivity to its (micro)environment. Phasor analysis yielded apparent phase lifetimes (τφ) of 3.70 ns in deionized water, 2.89 ns in imaging medium, 2.65 ns in 25% FBS, 2.99 ns in fresh human peritoneum, and 3.06 ns in rat tumor tissue (Figure 1f). The phase lifetime, derived from the angular position of the decay on the phasor plot, ^49^ therefore spanned 2.65–3.70 ns, a range of 1.05 ns. These values are not directly comparable to the 4.2-4.4 ns obtained by time-correlated single photon counting (TCSPC) on bulk dispersions because the two measurements differ in instrumentation, excitation repetition rate, and analysis. The value in water, 3.70 ns, was consistent with the 3.71 ns reported independently for the same particles.^45^ The environmental variation indicates that the RL-100 matrix does not completely shield aza-BODIPY from its surrounding environment. This observation is consistent with previous reports of shorter phase lifetimes for aza-BODIPY-labeled RL-100 NPs in serum and intestinal organoids than in water.^45^ The shorter lifetimes in peritoneum and tumor tissue likewise show that an aqueous reference alone cannot describe the tissue-associated signal. These matrix-specific phasor positions (τφ 2.65–3.70 ns, with 2.99 ns in fresh human peritoneum) therefore provide reference signatures for NP-associated fluorescence in tissue. Combined with its spectral separation from DOX, this establishes the tracer as both a carrier marker and a reporter of its local environment, forming the basis for ex vivo tissue imaging.

### 2.2. Spectral and Fluorescence Lifetime Characterization of NP Signals in Peritoneal Tissue

Tissue autofluorescence can complicate the detection of fluorescently labeled NPs when probe and endogenous emission overlap. In fresh human peritoneum, autofluorescence overlapped substantially with the emission of the amine-modified polystyrene FluoSpheres used in our previous EMDA study,^21^ with both showing maxima between approximately 540 and 560 nm (Figure 2a). The diBuO-aza-BODIPY emission (maximum at 718 nm) lies outside this window, 55 nm beyond the visible detection band (Figure 2a). After passive incubation with aza-BODIPY-labeled RL-100 NPs and washing, near-infrared fluorescence (excitation 698 nm; detection 703–750 nm) persisted along the exposed tissue surface (Figure 2b). Phasor analysis separated the fluorescence lifetime signatures of the tissue and NP-associated signals acquired in their respective spectral channels (Figure 2b). Lifetime contrast has previously discriminated near-infrared NP reporters with similar emission spectra in living cells,^50^ and phasor analysis has enabled multiplexed imaging of probes with overlapping spectra.^51^ Here, spectral separation already isolated the tracer from tissue autofluorescence, so lifetime analysis instead reported on the tracer in its tissue environment. In fresh peritoneum, the NP-associated population showed a mean τφ of 2.99 ns (range 2.95– 3.03 ns; Figures 1f and 2b), which defined the tissue-associated NP signature used as a reference in subsequent experiments. The shorter lifetime than in water (3.70 ns) is consistent with the environmental response previously reported for aza-BODIPY-labeled RL-100 NPs.^45^ Depth-resolved analysis was performed on cryosections cut perpendicular to the exposed tissue surface and fixed in methanol. NP-associated fluorescence was therefore characterized under the same preparation conditions (Figure 2c). Untreated fixed peritoneum showed low near-infrared background, whereas NP-treated tissue exhibited an NP-associated population with a mean τφ of 2.76 ns (range 2.69–3.03 ns), compared with 2.99 ns in fresh tissue (Figure 2b). The fixed-tissue phasor signature was used to select NP-associated fluorescence before depth-resolved intensity quantification. Together with the tissue-free NP reference and the corresponding phasor distributions, these controls supported assigning the detected near-infrared signal to NP-associated fluorescence. Tissue morphology and near-infrared fluorescence also remained suitable for imaging after fixation. Whereas previous characterization of this NP chemistry demonstrated its application in organoids,^45^ the present observations support its use in methanol-fixed peritoneal sections, with reference signals established for this preparation. Together, spectral detection, lifetime characterization, and untreated-tissue controls provided the basis for assessing NP distribution after passive delivery and EMDA.

**Figure 2.**
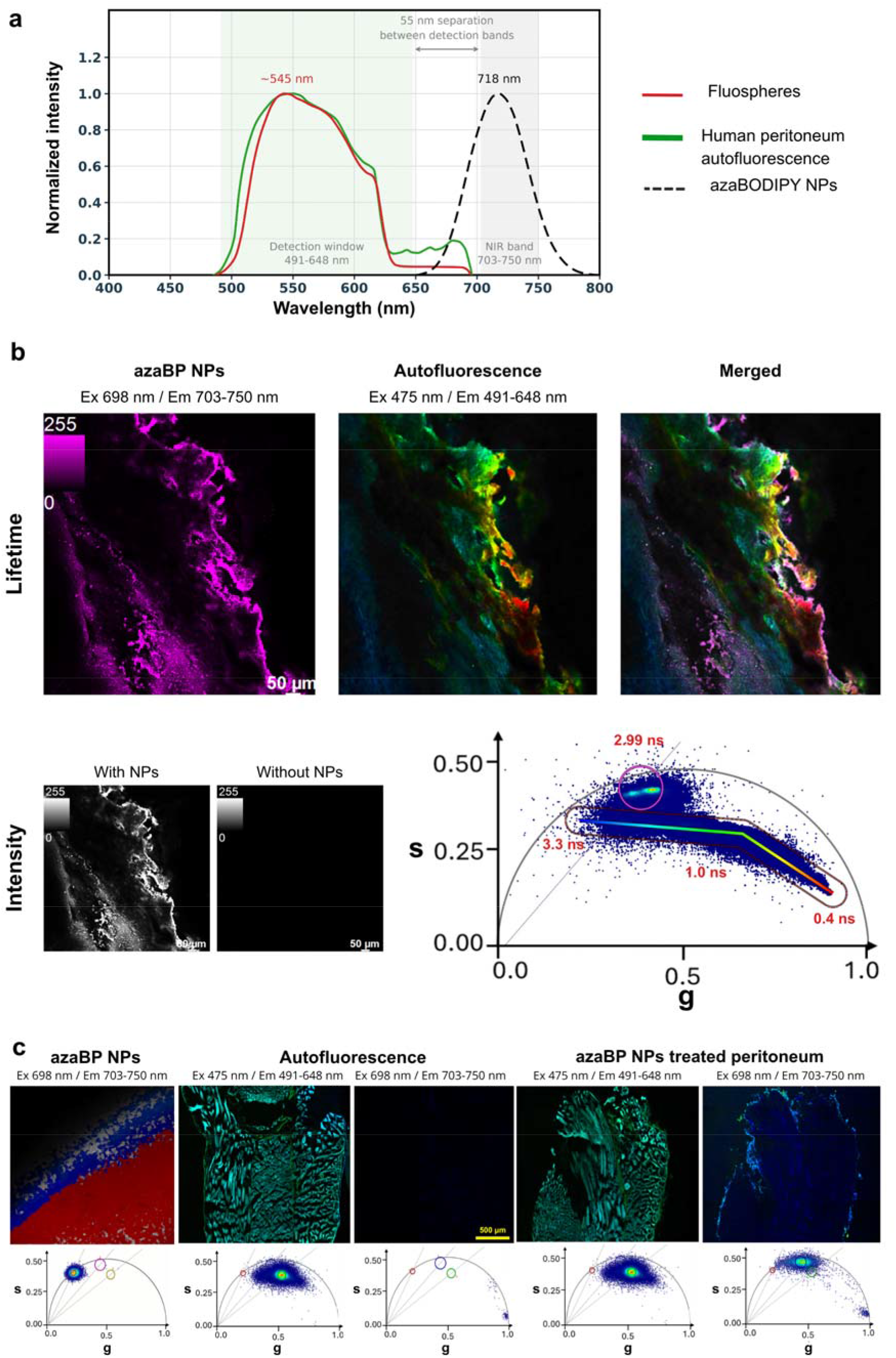
Spectral and fluorescence lifetime characterization of NP-associated signals in fresh and fixed human peritoneum. a) Normalized emission spectra of amine-modified polystyrene FluoSpheres (red), untreated human peritoneum (green), and diBuO-aza-BODIPY-labeled RL-100 NPs (black dashed line; λexc = 635 nm). Shaded regions indicate the visible (491–648 nm) and NIR (703–750 nm) detection bands. b) Fresh peritoneum following passive NP incubation and washing, imaged without fixation. Top: phasor-selected NP fluorescence, the corresponding NIR intensity image, and an untreated NIR control. Grayscale images use identical intensity limits. Bottom: autofluorescence, merge, and phasor plot. Autofluorescence is pseudocolored according to the phasor trajectory, with selected lifetimes annotated; the magenta phasor mask identifies the NP-associated population (mean τφ = 2.99 ns). Scale bars: 50 µm. c) Tissue-free NPs in water, untreated methanol-fixed peritoneum, and NP-treated methanol-fixed peritoneum, with corresponding phasor plots below. Tissue samples are shown separately in the visible and NIR channels, with identical intensity limits for untreated and treated images within each channel. Circular phasor masks indicate the tissue-free NP reference (red; τφ = 3.70 ns), tissue autofluorescence (green; τφ = 2.45 ns), and NP-associated fluorescence in fixed peritoneum (blue; mean τφ = 2.76 ns, range 2.69–3.03 ns). The fixed-tissue NP signature served as the reference for depth analysis. Scale bars: 500 µm. Visible channel: λexc = 475 nm, λem = 491–648 nm; NIR channel: λexc = 698 nm, λem = 703–750 nm. Fresh-tissue observations were obtained from five specimens.

### 2.3. EMDA-Enhanced NP Penetration in Ex Vivo Fixed Human Peritoneal Tissues

NP penetration into ex vivo human peritoneum was compared under passive diffusion (PD) and EMDA at three NP concentrations. A circular phasor mask centered at τφ = 2.76 ns, with the radius parameter set to 19 in LAS X, was first applied to select NP-associated pixels (Figure 3a). Mean near-infrared intensity was then quantified in three contiguous 100 µm ROIs extending from the exposed mesothelial surface (Figure 3b). Mean NP-associated fluorescence intensity in the surface-adjacent region (ROI 1, 0–100 µm) did not differ significantly between PD and EMDA at any tested concentration. However, the groups diverged in depth. EMDA produced significantly higher intensity in ROI 2 and ROI 3 at every concentration (p < 0.05 at 73 and 29.2 µg mL□¹; p < 0.001 at 14.6 µg mL□¹). Under PD, NP-associated signal remained largely confined to the first 100 µm, whereas under EMDA it was detectable throughout the 300 µm analyzed. Thus, the external electric field significantly enhanced NP penetration in healthy human peritoneum, consistent with our earlier observations.^21^ Mechanistically, electromigration of charged carriers and electroosmotic transport provide plausible explanations for the deeper penetration of cationic RL-100 NPs^18,48^ although field-induced changes in tissue permeability cannot be ruled out as an additional mechanism.

**Figure 3.**
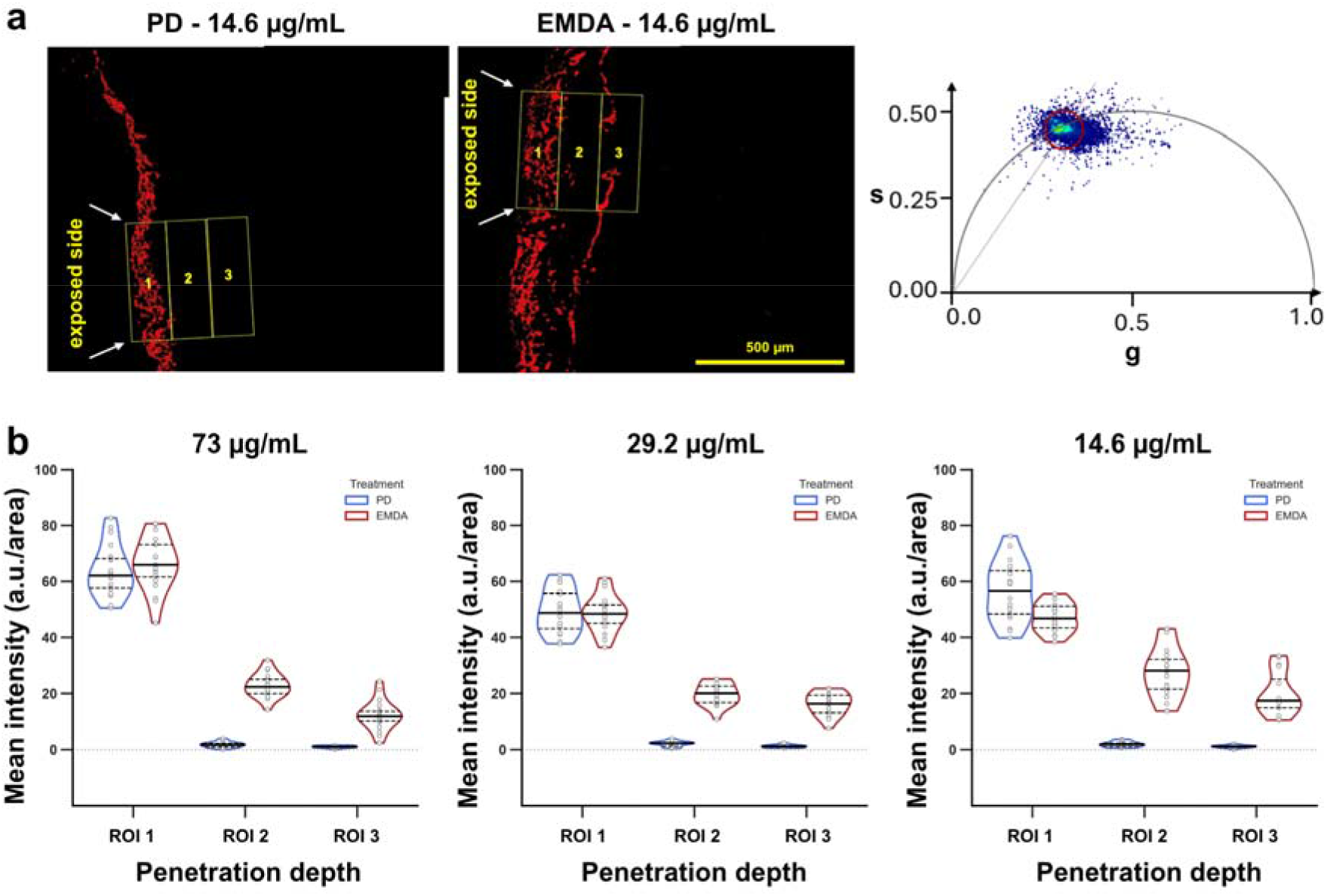
EMDA-enhanced NP penetration in ex vivo human peritoneum. a) Representative near-infrared images after PD or EMDA at 14.6 µg mL□¹, displayed with identical intensity limits and with the quantified ROIs overlaid. Arrows indicate the exposed mesothelial surface. The phasor plot shows the circular phasor mask (center τφ = 2.76 ns; radius parameter 19 in LAS X) used to select NP-associated fluorescence before quantification. Scale bars: 500 µm. b) Mean near-infrared fluorescence intensity by depth after PD or EMDA at azaBP-NP concentrations of 73, 29.2, and 14.6 µg mL ¹, quantified in three consecutive 100 µm ROIs from the exposed mesothelial surface (ROI 1, 0–100 µm; ROI 2, 101–200 µm; ROI 3, 201–300 µm). Violin plots show the distribution of individual ROI measurements, with medians (solid lines) and quartiles (dashed lines). The dataset comprised 12 tissue samples from three independent experiments. Comparisons used two-way repeated-measures ANOVA with Tukey-adjusted post hoc tests.

Enhanced penetration was observed across all three concentrations, and deep-ROI intensity under EMDA did not increase with NP concentration over the range tested, indicating a concentration-independent effect within this range (Figure 3b). The lowest concentration, 14.6 µg mL□¹, provided detectable signal throughout the analyzed depth under EMDA and was therefore selected for the subsequent healthy-rat experiments; in the tumor study, NPs were dosed by DOX equivalent (Section 2.7). Lifetime information was used to segment the NP-associated pixels (valid τφ range 2.69–3.03 ns), whereas fluorescence intensity within these pixels provided the quantitative measure of spatial distribution. Because penetration was measured on tissue cross-sections, depth represents distance from the exposed mesothelial surface rather than optical imaging depth. The profiles establish deeper NP-associated fluorescence under EMDA within the 300 µm range examined but do not define the maximum penetration depth. Having demonstrated enhanced NP penetration in ex vivo human tissue, we next examined its distribution in the rat peritoneum in vivo.

### 2.4. EMDA-Enhanced NP Penetration in Rat Peritoneum In Vivo

To test whether the effect persisted in vivo, aza-BODIPY-labeled RL-100 NPs without DOX were administered intraperitoneally to healthy rats at 14.6 µg mL ¹ using PD or EMDA, with applied currents up to 1.6 mA. Heart rate, respiratory rate, and oxygen saturation remained within the physiological range and were comparable between groups (Table S1). Procedural controls receiving current without NPs showed no detectable near-infrared signal above background under the same imaging conditions (Figure S3).

In methanol-fixed rat peritoneum, NP-associated fluorescence formed a distinct phasor population with a mean τφ of 2.76 ns (range 2.69–3.03 ns), matching the signature observed in methanol-fixed human peritoneum and shorter than the tissue-free NP reference in water (3.70 ns; Figure 4a, b). After phasor selection, mean fluorescence intensity was quantified in successive 100 µm ROIs extending from the exposed mesothelial surface. Under PD, NP-associated fluorescence was predominantly concentrated in the outermost 100 µm (ROI 1). EMDA produced significantly higher intensity in ROI 2 (101–200 µm) and ROI 3 (201–300 µm) in both right and left parietal peritoneum (p < 0.001; Figure 4c, d). Intensity declined with depth under both conditions, yet EMDA increased NP-associated fluorescence in the deeper regions throughout the analyzed range, confirming earlier observations using polystyrene NPs.^21^ These observations suggest that electrically enhanced NP transport persists in living rat abdominal wall tissue, where the vascular system can strongly influence the pharmacokinetics of IP-delivered drugs. The shorter lifetime of the tissue-associated tracer relative to tissue-free NPs is consistent with the environmental sensitivity reported for the same aza-BODIPY-labeled RL-100 platform.^45^ In this experiment, lifetime identified the NP-associated fluorescence population, whereas intensity quantified its spatial distribution. The lifetime shift therefore indicates that the tracer experienced a different local environment in tissue, rather than as a measure of penetration or proof that the NP matrix remained intact.

**Figure 4.**
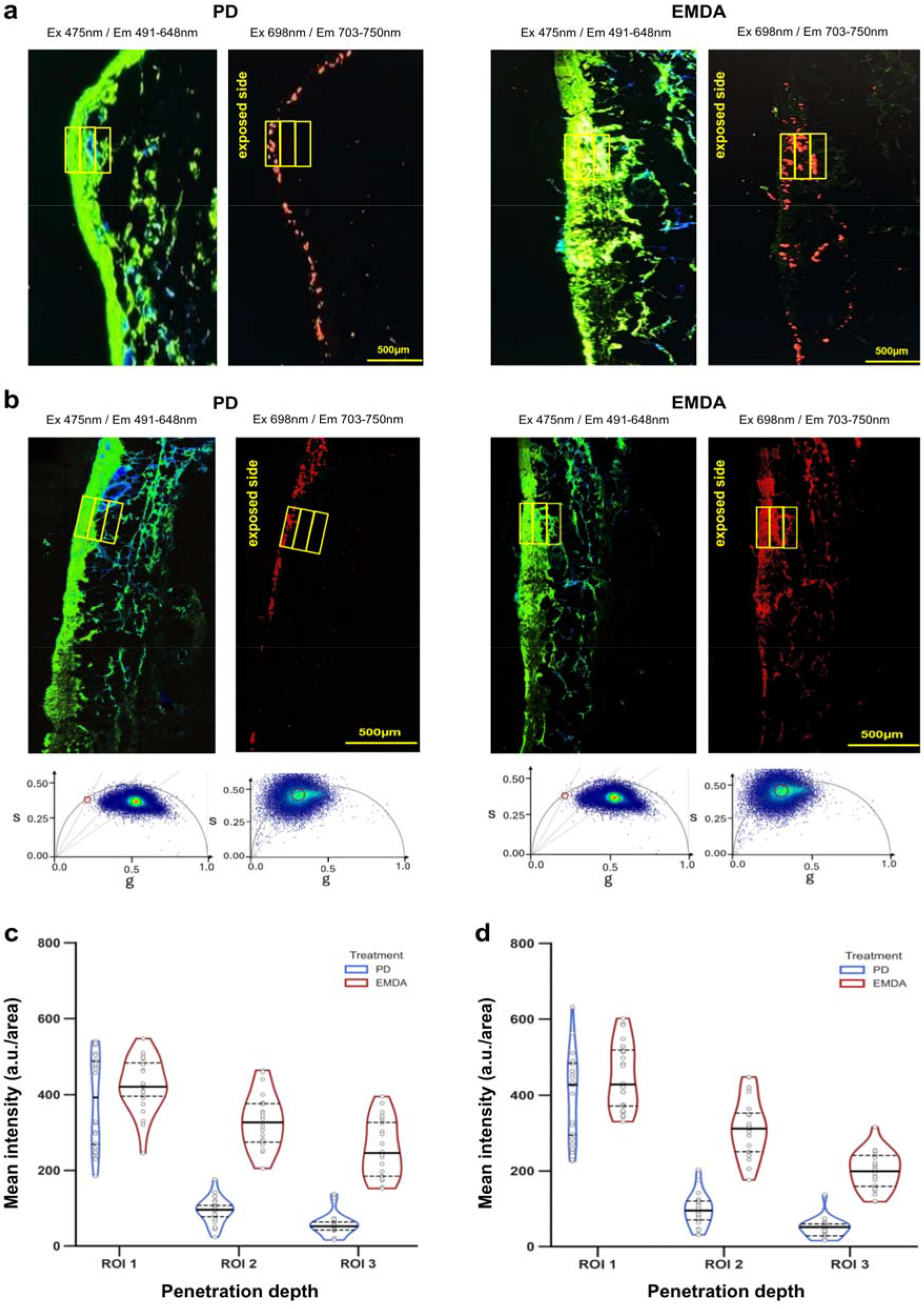
Depth-resolved NP distribution in rat parietal peritoneum following PD or EMDA. a,b) Representative images of a) right and b) left parietal peritoneum after intraperitoneal administration of azaBP NPs at 14.6 µg mL□¹ under passive diffusion (PD) or electromotive drug administration (EMDA). For each condition, images show tissue autofluorescence (λexc = 475 nm; λem = 491–648 nm) and the near-infrared NP signal (λexc = 698 nm; λem = 703–750 nm). Arrows indicate the exposed mesothelial surface, and boxes mark the three ROIs used for quantification. Phasor plots corresponding to the two detection channels are shown below the left-peritoneum images in panel b. Circular phasor masks mark the tissue-free NP reference (red; τφ = 3.70 ns) and NP-associated population (center τφ 2.76 ns, radius parameter 19) selected before intensity quantification. c,d) Mean near-infrared fluorescence intensity by depth in c) right and d) left parietal peritoneum. ROI 1, ROI 2, and ROI 3 represent distances of 0–100, 101–200, and 201–300 µm, respectively, from the exposed mesothelial surface. Violin plots show the distribution of individual ROI measurements, with medians (solid lines) and quartiles (dashed lines); n = 3 animals per group. Comparisons were performed using two-way repeated-measures ANOVA with Tukey adjustment. Scale bars: 500 µm.

### 2.5. Intracellular DOX Release and Nuclear Delivery from RL-100 NPs in Colorectal Carcinoma Cells

Having established the extent of RL-100 NP penetration using PD and EMDA, we next examined whether the carrier released DOX after cellular uptake, how the released drug progressed toward its nuclear target, and which of two DOX-equivalent concentrations provided a sufficiently robust signal to evaluate intracellular release.

DOX fluorescence lifetime is sensitive to its molecular environment, enabling FLIM to complement spatial imaging when assessing carrier-associated and intracellular drug states. Previous studies have used lifetime differences to investigate intracellular DOX release and changes in its interactions with nuclear components.^27,52^ In our cell-free measurements, DOX encapsulated in RL-100 NPs exhibited a phase lifetime of approximately 1.7 ns, compared with 1.1 ns for free DOX in imaging medium. These measurements provided formulation-specific references for interpreting intracellular fluorescence. Following nuclear entry, further lifetime changes can reflect interactions with DNA and changes in the chromatin environment; NP-delivered DOX was therefore evaluated against matched free-DOX controls and the separately detected carrier-associated signal.

Rat colorectal cancer cells (CC531s.2) were exposed to DOX/aza-BODIPY co-loaded RL-100 NPs (DOX azaBP NPs) or free DOX at 2 and 20 µM DOX-equivalent concentrations and imaged after 1, 5, and 24 h. These cell-free reference lifetimes guided the interpretation of the subsequent intracellular trajectory rather than serving as fixed lifetime values for DOX in cells.

The intracellular distribution of DOX differed between the two delivery routes at the first time point (Figure 5a, Figure S4). At 1 h, NP-delivered DOX appeared predominantly as punctate cytoplasmic fluorescence, whereas free DOX showed more extensive nuclear accumulation. Nuclear DOX was nevertheless detectable after NP delivery, indicating that some payload had already separated from the carrier. At 5 h, the punctate cytoplasmic pattern diminished, and NP-delivered DOX progressively accumulated in the nuclei. By 24 h, nuclear DOX was evident following both delivery routes, whereas the aza-BODIPY signal remained predominantly cytoplasmic, indicating spatial separation of the carrier-associated tracer from DOX that had reached the nucleus (Figure S5). Phasor analysis of DOX lifetime at 1 h showed that NP-delivered DOX occupied the longer-lifetime phasor masks, consistent with predominantly carrier-associated drug, whereas free DOX occupied the shorter-lifetime phasor masks, consistent with released drug interacting with nuclear components (Figure 5a). Dye-only NPs and untreated cells produced no detectable signal above background in the DOX channel under the same acquisition conditions (Figure S5). This intracellular redistribution contrasts with the limited leakage observed during prolonged storage of concentrated NP dispersions in water. The lower NP concentration and different biological environment during cellular exposure may favor DOX partitioning out of the polymer matrix, although the relative contributions of dilution and interactions with medium components or cellular membranes were not determined.

**Figure 5.**
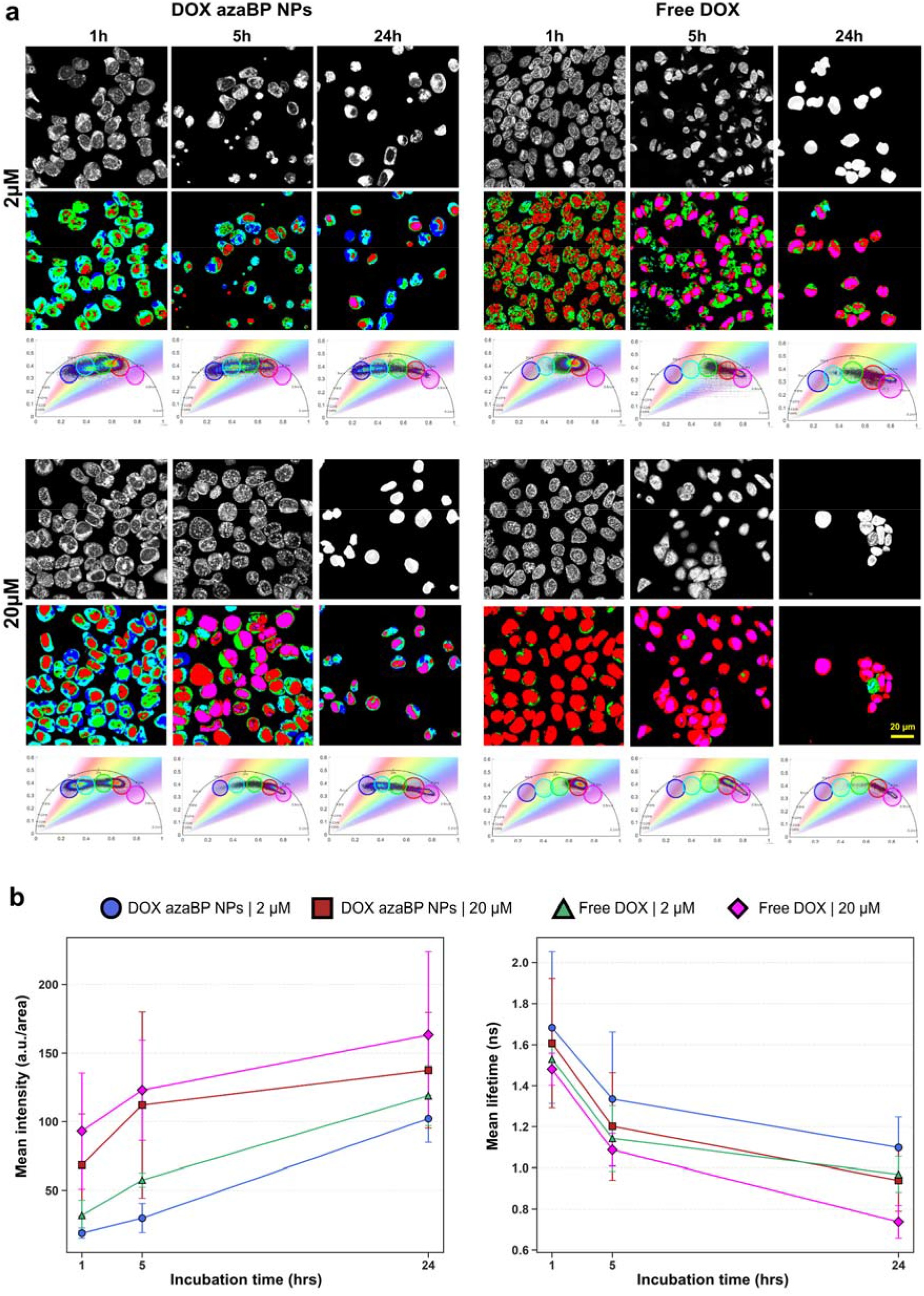
Intracellular DOX release and nuclear delivery from RL-100 NPs in CC531s.2 cells. a) Representative images of cells exposed to DOX-azaBP-NPs or free DOX at DOX-equivalent concentrations of 2 and 20 µM for 1, 5, or 24 h. For each condition, rows show DOX fluorescence intensity in grayscale, pixels pseudo colored by five circular phasor masks (blue to magenta, decreasing phase lifetime), and the corresponding phasor plot. The complete image set, including phase-lifetime-coded images, is shown in Figure S4. b) Mean DOX fluorescence intensity (left) and mean DOX phase lifetime (right) within segmented nuclear regions. Data are presented as mean ± SD; at least 10 nuclei per field were analyzed in each of three independent experiments. Lifetime values are reported descriptively. Scale bar: 20 µm.

Quantification was restricted to segmented nuclear regions. Nuclear DOX intensity increased with time under all conditions and was higher at 20 µM than at 2 µM (Figure 5b). At 2 µM, the nuclear intensity of NP-delivered DOX remained relatively low at 1 and 5 h and increased more markedly by 24 h. At 20 µM, nuclear DOX was prominently detected at earlier time points and remained more intense throughout the experiment.

Nuclear phase lifetime decreased progressively from 1 to 24 h at both concentrations (Figure 5b). At 2 µM, it decreased from 1.68 to 1.10 ns for NP-delivered DOX and from 1.55 to 0.97 ns for free DOX. At 20 µM, it decreased from 1.61 to 0.94 ns for NP-delivered DOX and from 1.48 to 0.73 ns for free DOX.

Since the analysis was restricted to nuclear regions, the measured lifetime reflects the state of nuclear DOX rather than the amount of DOX remaining associated with NPs in the cytoplasm. Free DOX also showed a progressive lifetime decrease despite the absence of a carrier, demonstrating that intracellular processing contributes to the trajectory. Nuclear DOX lifetime is sensitive to its interaction with DNA and to changes in the chromatin microenvironment during the early stages of apoptosis.^52^ The progressive decrease in lifetime could therefore reflect continued interaction of released DOX with nuclear components, although lifetime alone cannot identify a specific binding state or demonstrate apoptosis without corresponding biological markers.

The matched free-DOX conditions provided a controlled reference for interpreting NP-delivered DOX. At every concentration and time point, the mean nuclear lifetime of NP-delivered DOX was longer than that of free DOX. This consistent pattern suggests that NP-delivered DOX reached a different or earlier nuclear state, compatible with delayed intracellular availability following carrier-mediated delivery. The independent carrier channel supports this interpretation: the aza-BODIPY-labeled carrier remained cytoplasmic while DOX reached the nucleus (Figure S5), so nuclear DOX following NP treatment represents released payload rather than intact NP cargo. Spatial separation therefore supports intracellular payload release, while nuclear lifetime reports the subsequent state of the released drug relative to fully available free DOX.

Previous FLIM studies have distinguished carrier-associated from released DOX and followed its movement from the cytoplasm to the nucleus.^27,31,35^ An intracellular DOX lifetime of approximately 1.15 ns has also been reported following delivery from PLGA NPs, supporting the assignment of the shorter-lifetime population to intracellularly available drug.^32^ The present study adds a matched free-DOX time course at two concentrations, allowing the lifetime of NP-delivered DOX to be interpreted against drug of known availability under the same cellular conditions.

Both concentrations showed increasing nuclear DOX intensity and decreasing phase lifetime over time, but the magnitude and timing of the response differed. At 2 µM, nuclear accumulation of NP-delivered DOX was limited at 1 and 5 h and became evident only at 24 h. The 20 µM condition therefore provided the clearest readout of intracellular release and was selected for cell-based release assessment. Notably, the nuclear lifetime of NP-delivered DOX at 20 µM closely followed that of free DOX at 2 µM (1.61 vs 1.55 ns at 1 h; 0.94 vs 0.97 ns at 24 h), suggesting that the DOX released from NPs reached a nuclear state comparable to that produced by approximately 2 µM free drug. Lifetime kinetics therefore allowed a semi-quantitative estimate of the effective released dose, which nuclear intensity could not provide because of its larger variability (Figure 5b). This selection does not translate directly to the in vivo dose. Overall, the results demonstrate concentration- and time-dependent release of DOX from RL-100 NPs, followed by nuclear accumulation and changes in the functional state of the released drug.

### 2.6. Tumor-Associated Retention of Aza-BODIPY-Labeled RL-100 NPs Revealed by Whole-Animal and Ex Vivo Imaging

Whole-animal near-infrared imaging was used to examine the distribution and retention of azaBP NPs in tumor-bearing rats (Figure 6a). Acquisition settings were selected for the near-infrared tracer; DOX was not evaluated because its maximum emission near 590 nm lies outside the Cy5.5 detection window. The measurements therefore describe the aza-BODIPY-associated signal and should not be interpreted as whole-animal DOX biodistribution.

**Figure 6.**
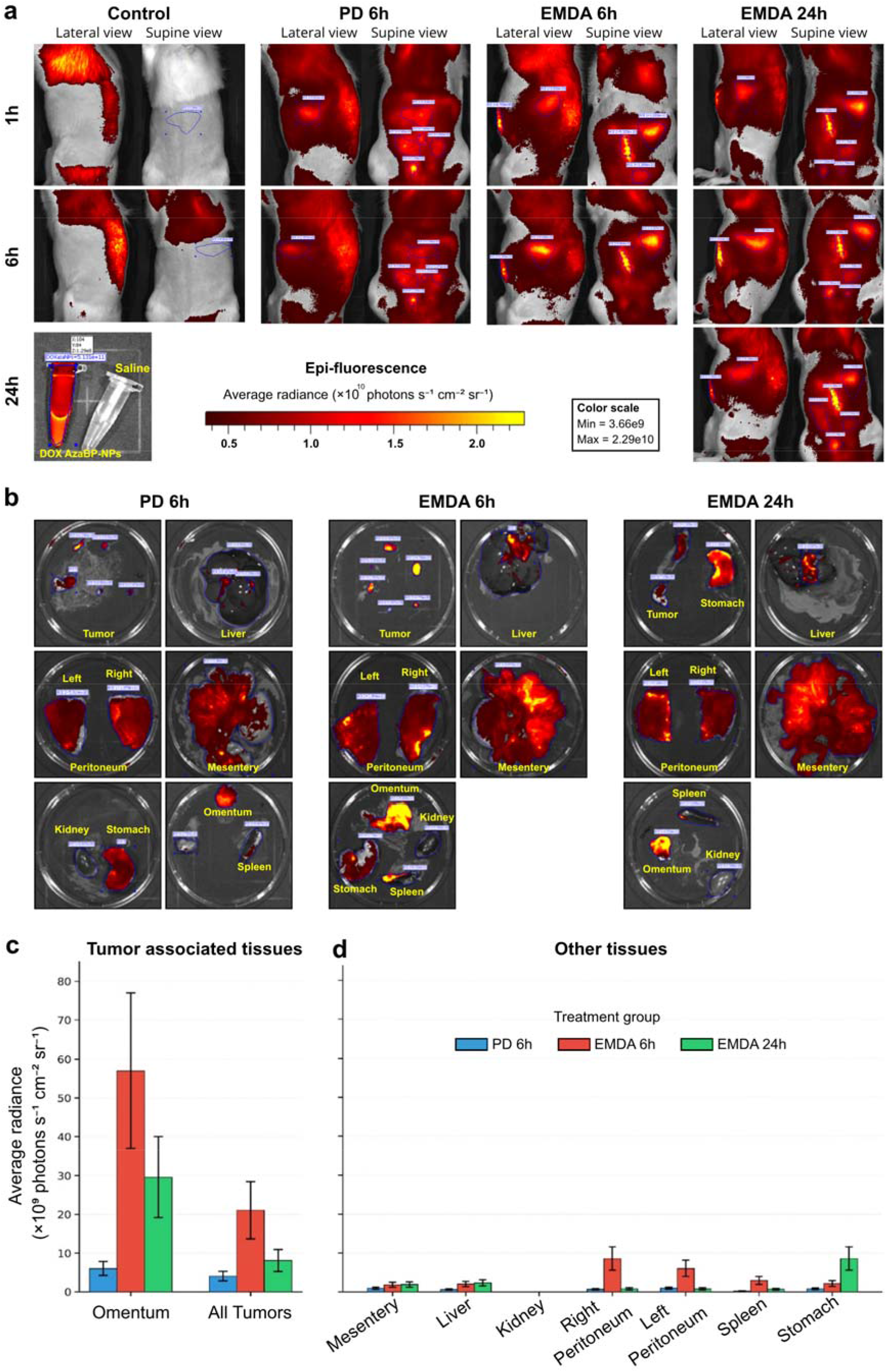
Whole-animal and ex vivo distribution of aza-BODIPY-labeled RL-100 NPs in tumor-bearing rats. a) Representative lateral and supine near-infrared images of a saline-treated control and animals receiving DOX azaBP NPs by passive diffusion (PD) or electromotive drug administration (EMDA). Columns identify the treatment group and its endpoint; rows give the imaging time point, so animals assigned to the 6 h endpoint were imaged at 1 and 6 h and those assigned to the 24 h endpoint at 1, 6 and 24 h. Images are displayed using a common radiance scale (minimum 3.66 × 10□, maximum 2.29 × 10¹□ photons s□¹ cm□² sr□¹) and outlined regions indicate the regions of interest used for quantification. The inset shows tubes of saline and of the DOX azaBP NPs dispersion imaged under identical settings. b) Representative ex vivo images of excised tumor nodules, tumor-bearing omentum, right and left parietal peritoneum, mesentery, liver, kidney, spleen and stomach, harvested 6 h after PD, 6 h after EMDA, or 24 h after EMDA; tissues were grouped in dishes for imaging, and the arrangement differs between panels. c,d) Average radiance in c) tumor-associated tissues, comprising tumor-bearing omentum and all excised tumor nodules, and d) other tissues, for PD at 6 h (blue), EMDA at 6 h (red), and EMDA at 24 h (green). Bars show group means and error bars indicate SD; n = 3 animals per group. No statistical comparison was performed; group differences are descriptive.

Background-subtracted fluorescence was predominantly detected within the abdomen of treated animals, with no signal above control levels in the thorax or extremities. At the matched 6 h endpoint, abdominal fluorescence was stronger and more localized after EMDA compared to PD. High-intensity regions corresponded with tumor tissue locations identified at necropsy. Signal remained detectable 24 h after EMDA-mediated delivery, although its distribution and intensity differed from those observed at 6 h. A focal dorsal signal in one control animal was attributed to fur autofluorescence and excluded from the quantitative regions of interest.

Ex vivo imaging enabled direct comparison of the excised tissue (Figure 6b–d). At 6 h, average radiance in tumor nodules was approximately 21 × 10□ photons s□¹ cm□² sr□¹ after EMDA and 4 × 10□ photons s□¹ cm□² sr□¹ after PD, representing an approximately fivefold difference. The highest signal was measured in the tumor-bearing omentum, reaching approximately 57 × 10 photons s□¹ cm□² sr□¹ after EMDA compared with 6 × 10□ photons s□¹ cm□² sr□¹ after PD. Right and left parietal peritoneum showed the same directional difference, with values of approximately 8.6 and 6.0 × 10□ photons s□¹ cm□² sr□¹ after EMDA compared with 0.7 and 1.1 × 10□ photons s□¹ cm□² sr□¹ after PD.

Liver, kidney, spleen, and bowel mesentery remained below 3 × 10□ photons s□¹ cm□² sr□¹ in all groups. The resulting distribution is consistent with preferential retention in tumor tissue and directly exposed peritoneal tissue. At 24 h after EMDA, radiance decreased to approximately 8 × 10□ photons s□¹ cm□² sr□¹ in tumor nodules and 30 × 10□ photons s□¹ cm□² sr□¹ in the omentum but remained detectable at both sites. Because 6 h was the primary time point for comparing delivery methods, the 24 h time point was examined only after EMDA to assess signal persistence; these values therefore describe persistence under the EMDA condition rather than a difference between delivery methods. Gastric signal increased to approximately 8.6 × 10□ photons s□¹ cm□² sr□¹ at 24 h, indicating redistribution of the tracer within the abdominal cavity. Gastrointestinal clearance cannot be inferred because luminal localization and excretion were not measured. Of note, aza-BODIPY fluorescence identifies the tracer but does not prove that the complete NP is preserved after administration. Redistribution of fluorescent labels can cause the optical signal to diverge from nanocarrier fate, prompting the development of aggregation-caused-quenching tracers for in vivo NP imaging.^53,54^ The strong retention of the hydrophobic aza-BODIPY dye during formulation characterization reduces this concern but does not eliminate it in vivo. Near-infrared NP probes have previously been developed for the detection and fluorescence-guided resection of peritoneal tumor deposits.^55–58^ The tumor-associated contrast and persistence observed here support further evaluation of the RL-100 formulation for image-guided identification of peritoneal disease.

### 2.7. Electromotive Delivery Extends Doxorubicin Penetration into Peritoneal Tumor Nodules

The intratumoral distribution of free and NP-delivered DOX was evaluated in a syngeneic rat model of colorectal PM. Tumor nodules developed following IP inoculation of 2.0 × 10□CC531s.2 cells and were treated on day 15, with tissue harvested after 6 or 24 h (Figure 7a). Tumor nodules identified at necropsy were confirmed histologically (Figure 7b, c). Free DOX and DOX azaBP NPs were administered at a DOX-equivalent dose of 3 mg kg□¹ using PD or EMDA. Heart rate, respiratory rate, and oxygen saturation remained within the physiological range and were comparable between groups during treatment (Figure S6b).

**Figure 7.**
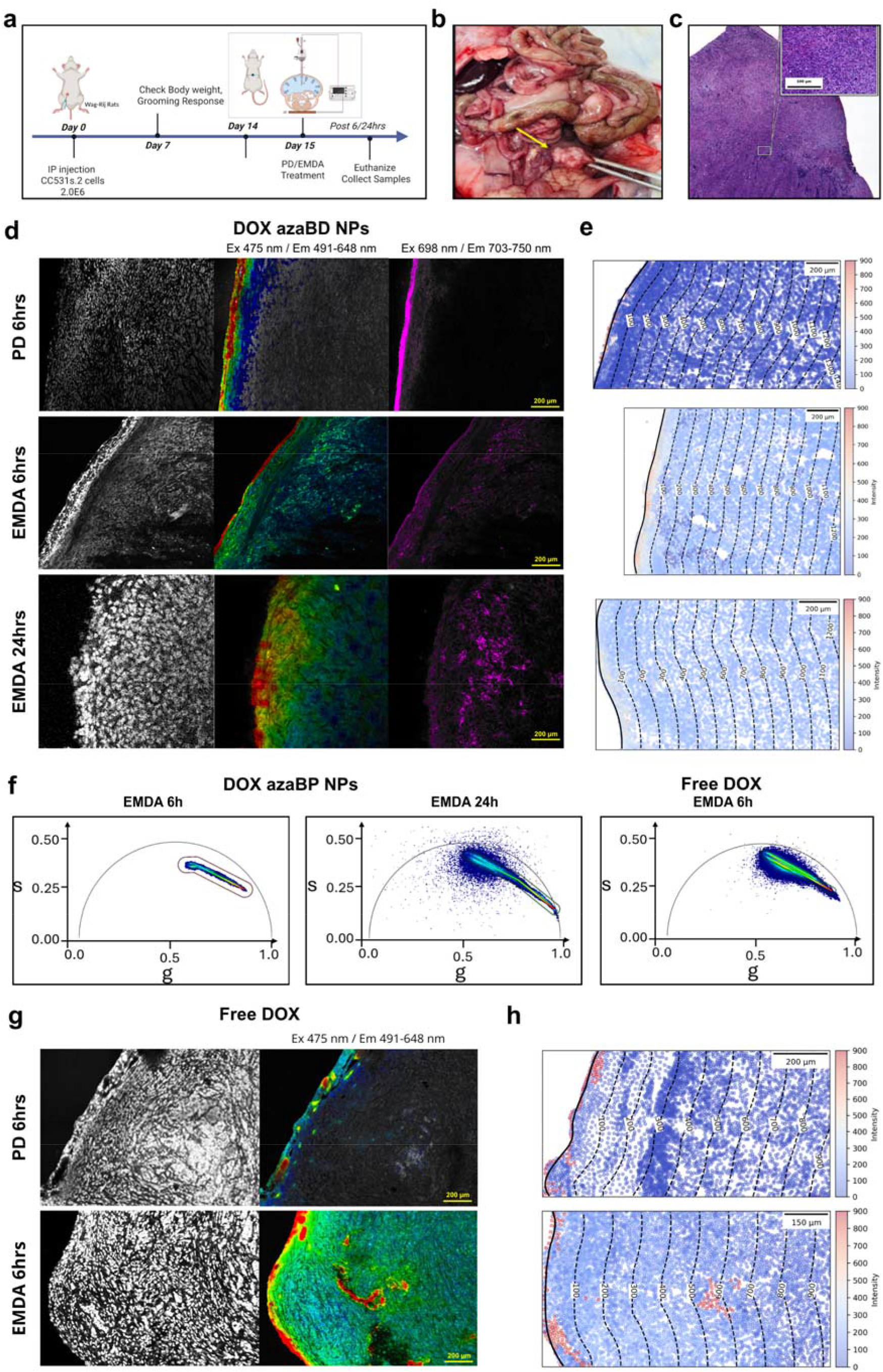
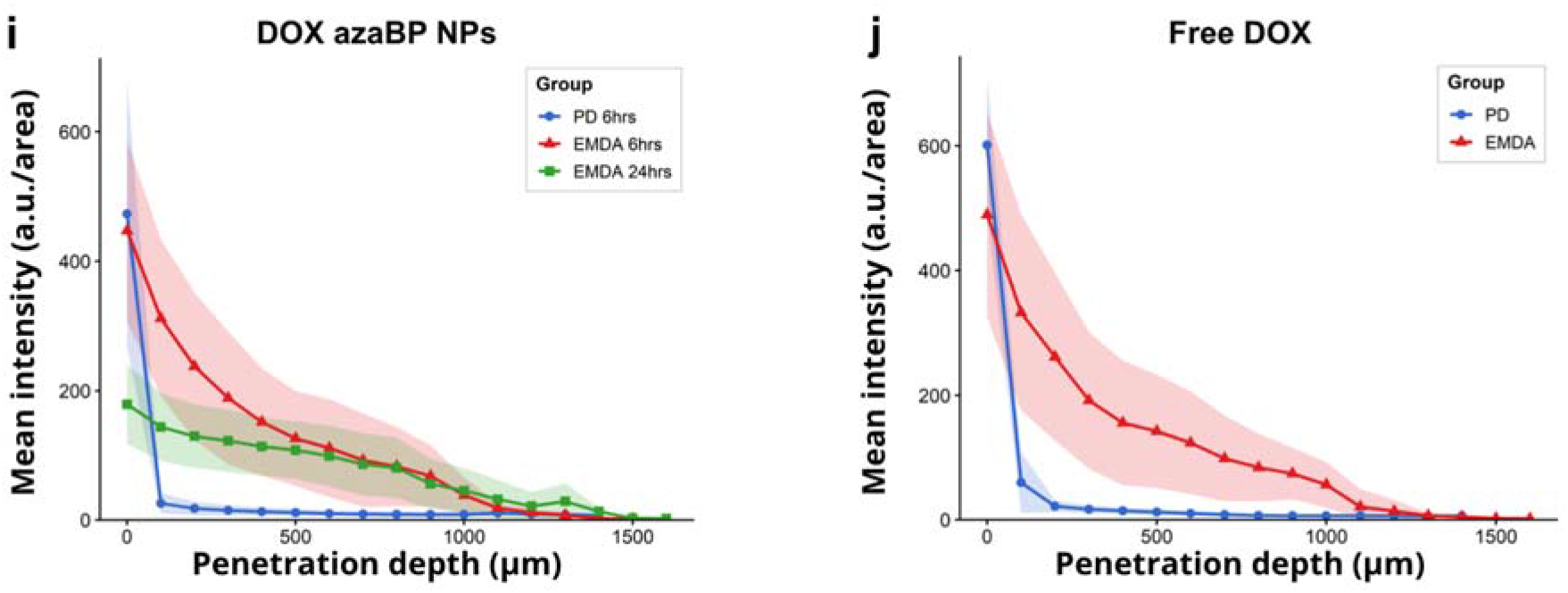
EMDA extends DOX penetration into colorectal peritoneal tumor nodules. a) Experimental timeline showing intraperitoneal inoculation of 2.0 × 10□ CC531s.2 cells on day 0, monitoring on days 7 and 14, PD or EMDA treatment on day 15, and tissue collection 6 or 24 h after treatment. b) Representative macroscopic view of peritoneal tumors at necropsy; the arrow indicates a tumor nodule. c) Representative hematoxylin and eosin-stained tumor section, with the boxed region enlarged in the inset. d) Representative tumor images following delivery of DOX azaBP NPs by PD at 6 h, EMDA at 6 h, or EMDA at 24 h. Columns show DOX-channel fluorescence intensity in grayscale, lifetime-coded DOX fluorescence (λexc = 475 nm; λem = 491–648 nm), and near-infrared aza-BODIPY fluorescence (λexc = 698 nm; λem = 703–750 nm). e) Corresponding DOX intensity maps with iso-depth contours at 100 µm intervals from the tumor surface. f) Representative DOX channel phasor plots following EMDA delivery of DOX azaBP NPs at 6 and 24 h and for free DOX at 6 h. g) Representative DOX intensity and lifetime-coded images following delivery of free DOX by PD or EMDA at 6 h. h) Corresponding DOX intensity maps with iso-depth contours at 100 µm intervals. Intensity maps in (e) and (h) share a common 0–900 intensity scale. i,j) Depth profiles of mean DOX fluorescence intensity following i) NP-delivered DOX and j) free DOX. Lines show the mean of the tissue-sample means at each depth, and shaded regions indicate ±SD across those means. Replicate measurements were averaged within each tissue sample. Three animals were included per group, with three tissue samples per animal. Profiles were generated from identically thresholded DOX intensity images without lifetime-based pixel selection. Separate generalized estimating-equation models with an AR(1) working correlation and a second-order polynomial function of depth were fitted for the NP and free-DOX groups. Scale bars are indicated within the panels.

Because DOX emission occupies the same spectral range as tissue autofluorescence, intratumoral DOX signal was interpreted relative to PD-treated nodules imaged under identical acquisition settings. Beyond the DOX front, intensity in PD-treated nodules dropped to near-background levels at the same threshold (Figure 7e, i), indicating that the deep signal after EMDA exceeded autofluorescence in comparable tumor tissue. The quantitative penetration profiles, however, were calculated from DOX intensity images using the same intensity threshold across all treatment groups and depths; they were not generated from lifetime-selected pixels.

Across all treatment groups, the mean DOX phase lifetime in tumor tissue was 1.39 ns, with group means ranging from 1.37 to 1.40 ns and individual values spanning approximately 0.8–1.7 ns (Figure 7f). These values were within the range measured for intracellular DOX in CC531s.2 cells and were shorter than the predominantly NP-associated state observed during early cellular exposure in Section 2.5. The similarity between the free-DOX and NP-delivered DOX groups is consistent with substantial payload release before tissue harvesting. DOX phase lifetime increased modestly with depth, from approximately 1.28 ns at the tumor surface to 1.42 ns at 700 µm. This shift may reflect differences in local DOX concentration, molecular interactions, or tumor microenvironment across the tissue depth. It was therefore treated descriptively and was not used to calculate penetration or release. The lifetime measurement identifies changes in DOX state and environment, whereas the intensity profiles describe its spatial distribution.

Delivery method strongly affected the distribution of DOX within tumor nodules (Figure 7d,e,g–j). Separate generalized estimating equation models were fitted for free DOX and NP-delivered DOX because the two formulations were not compared directly within a single statistical model. For free DOX, superficial intensity did not differ between PD and EMDA (Wald χ² = 0.04, p = 0.84), whereas the group-by-depth interaction was highly significant (χ² = 45, df = 2, p = 1.6 × 10□¹□). Depth was also a strong predictor of intensity (χ² = 1172, df = 2, p < 0.001). For NP-delivered DOX, EMDA at 6 h produced modestly higher superficial intensity than PD at 6 h (χ² = 6.9, p = 0.009), whereas superficial intensity was lower in the EMDA group examined at 24 h (χ² = 72.1, p < 0.001). The overall group-by-depth interaction was significant (χ² = 339, df = 4, p < 0.001), and depth remained a strong predictor of intensity (χ² = 524, df = 2, p < 0.001). These results show that EMDA changed both the initial NP-delivered DOX signal and its subsequent distribution with depth and time.

With PD, DOX was largely confined to the outer 100–200 µm of tumor tissue, and intensity decreased rapidly at greater depth. EMDA extended detectable DOX to approximately 1200 µm for both free and NP-delivered formulations. High-resolution mosaics of EMDA-treated nodules at 6 h further illustrate DOX distribution from the tumor surface into deeper tissue at cellular resolution (Figures S7 and S8). Free DOX produced higher intensity at 6 h, consistent with its immediate availability without a preceding release step.

The marked extension of DOX penetration is relevant because transport after IP administration is restricted by elevated IFP, extracellular-matrix resistance, and limited inward convection.^2,59^ Indeed, computational models of IP chemotherapy predict steep surface-to-core concentration gradients in tumor tissue, consistent with the profiles observed here under PD.^59^ For NP-delivered DOX, the measured depth profile may reflect movement of drug-loaded carriers, drug released during delivery, or both. Matrix architecture and interactions with charged matrix components can influence carrier transport and retention,^60,61^ but their contributions were not isolated here. Similar detectable penetration depths therefore do not imply identical transport mechanisms or equivalent delivery between formulations. The experiment demonstrates enhanced intratumoral transport under EMDA; it does not establish that the electric field alters the rate of DOX release from the NPs.

The NP formulation also showed a distinct temporal profile. At 24 h after EMDA, superficial intensity was lower than at 6 h, while DOX remained distributed across much of the analyzed depth and exceeded the 6 h profile beyond approximately 700 µm (Figure 7i). Read together with the gradual intracellular release observed in Section 2.5, this pattern is compatible with continued local availability of DOX from the carrier after the initial exposure period. Free IP drug is cleared relatively rapidly, limiting the duration of tumor contact after a single administration,^3,62^ whereas controlled-release intraperitoneal systems are designed to extend local exposure.^63,64^ DOX-loaded carriers have also produced improved therapeutic outcomes relative to free drug in preclinical PM models.^9,10^ The 24 h results therefore demonstrate persistence of NP-delivered DOX after EMDA but cannot establish whether this persistence resulted from the carrier, EMDA, or their combination.

This study has several limitations. Animal experiments included three animals per group, limiting precision and generalizability. The DOX-co-loaded formulation was not independently characterized for particle size and zeta potential, and the qualitative retention assay did not quantify encapsulation efficiency or release kinetics. Carrier-associated fluorescence does not establish NP integrity in vivo, while DOX lifetime and nuclear localization support intracellular release without quantifying the released fraction, intracellular concentration, or therapeutic activity. The 24 h assessment was restricted to EMDA-delivered NPs, preventing comparison of persistence with passive delivery or free DOX at that time point. Further studies are needed to determine whether the observed improvements in delivery translate into improved tumor control.

## 3. Conclusion

A dual-reporter NP formulation was developed by co-loading DOX and a near-infrared aza-BODIPY tracer within a cationic RL-100 matrix, allowing carrier-associated fluorescence and payload behavior to be evaluated separately in autofluorescent tissue. EMDA enhanced NP penetration in ex vivo human peritoneum and healthy rat peritoneum, and DOX penetration in a syngeneic model of colorectal peritoneal metastasis. In tumors, both free and NP-delivered DOX remained detectable to approximately 1200 µm after EMDA, compared with less than 200 µm after passive delivery. For free DOX, deeper penetration occurred without a significant difference in surface intensity, supporting enhanced tumor tissue transport. NP-delivered DOX remained detectable 24 h after EMDA.

Cellular imaging showed predominantly cytoplasmic carrier-associated fluorescence alongside progressive nuclear accumulation of DOX, supporting intracellular payload release. By linking organ-level carrier-associated retention, depth-resolved drug penetration, and intracellular release within one experimental system, this work extends the evaluation of EMDA beyond carrier distribution alone. The combination of electrically enhanced transport and separate monitoring of carrier and payload provides a preclinical basis for NP-assisted intraperitoneal delivery in PM.

## 4. Experimental Section

### Nanoparticle Preparation

Near-infrared (NIR) fluorescent polymeric NPs were prepared by nanoprecipitation from the cationic methacrylate copolymer Eudragit RL-100 (Evonik, Germany) doped with dibutoxy-aza-BODIPY (diBuO-aza-BODIPY), following previously described procedures.^45,47^ For dye-only particles, Eudragit RL-100 (160 mg) and diBuO-aza-BODIPY (1.6 mg, 1 *w/w*%) were dissolved in acetone (80 mL). Under vigorous stirring, deionized water (400 mL) was added within several seconds. Acetone and part of the water were removed under reduced pressure at 40 °C using a rotary evaporator, yielding approximately 20 mL of colloidal dispersion. The dispersion was filtered through a cotton plug to remove aggregates formed during evaporation. Glycerol was added to 25 w/w% in the final dispersion for storage stability during transport between the preparing and imaging laboratories, giving a stock concentration of 7.3 mg mL ¹. These particles are referred to as azaBP NPs.

For drug delivery and release studies, doxorubicin (DOX)-loaded aza-BODIPY co-loaded RL-100 NPs (DOX azaBP NPs) were prepared by combining DOX hydrochloride (16 mg, 10 *w/w*% with respect to polymer) dissolved in deionized water (9 mL) with the acetone solution of RL-100 and diBuO-aza-BODIPY prior to nanoprecipitation. The final acetone-to-water ratio was 9:1 (*v/v*). Precipitation and purification were performed as described above, yielding a stock dispersion at 5.4 mg mL□¹. NPs were prepared at Graz University of Technology and shipped to Ghent University for all imaging and animal experiments. All dispersions were stored at 4 °C, protected from light, and vortexed immediately before use.

### Nanoparticle Characterization

UV-vis absorption spectra were recorded on a Cary 50 UV-vis spectrophotometer (Varian, USA). Fluorescence emission and excitation spectra were recorded on a Fluorolog-3 luminescence spectrometer equipped with an NIR-sensitive R2658 photomultiplier (Hamamatsu, Japan) in a screw-capped quartz cuvette (Hellma, Germany). Fluorescence decays were acquired by time-correlated single-photon counting (TCSPC) on the same spectrometer using a NanoLED excitation source (λ = 635 nm) and a DeltaHub module (Horiba, Japan).

The diBuO-aza-BODIPY emission spectrum shown in Figure 2a was recorded on this instrument. The emission spectra of untreated human peritoneum and of the amine-modified polystyrene FluoSpheres shown alongside it were replotted from data acquired in our previous EMDA study,^21^ in which they were recorded on a separate confocal system; the wavelength range shown reflects the detection settings of that instrument. All three traces are normalized to their own maxima and are compared for spectral position only.

The azaBP NPs used in this study originated from the same preparation batch as those previously characterized by transmission electron microscopy (TEM) by Okkelman et al.^45^ Sample preparation, image acquisition, and particle-size analysis were performed as described therein. In this study, we determined particle diameter from representative TEM images of 150 individual NPs. Size was not determined separately for the DOX azaBP co-loaded formulation.

Hydrodynamic size and ζ-potential could not be determined directly by dynamic light scattering (DLS; Zetasizer, Malvern Panalytical, UK), because diBuO-aza-BODIPY absorbs the 633 nm instrument laser almost quantitatively. Size and charge were therefore taken from previous characterization of RL-100 NPs prepared by the identical procedure, comprising DLS of undoped batches and of batches doped with dyes transparent in the red region.^44^

Payload incorporation and retention were assessed by salt-induced precipitation: saturated sodium chloride was added to a concentrated dispersion (approximately 5 mg □¹) at 1:5 *v/v*. This procedure precipitated the NPs, leaving any released payload in the supernatant. Dispersions were examined immediately after preparation and after 18 months of storage at 4 °C protected from light.

### Tissue Sources, Animals, and Ethics

Fresh human peritoneal tissue was obtained from patients undergoing elective abdominal surgery at Ghent University Hospital following written informed consent, under approval from the Medical Ethics Committee of Ghent University Hospital (ONZ-2022-0548; Belgian registry B6702023000012) and in accordance with the Declaration of Helsinki. Immediately after excision, specimens were transferred into imaging medium (Dulbecco’s modified Eagle medium without phenol red, supplemented with 10 × 10□³ M glucose, 1 × 10□³ M pyruvate, 2 × 10□³ M glutamine, and 10 × 10□³ M HEPES; Thermo Fisher Scientific, USA) to preserve viability and native autofluorescence. Tissue sections of 100 µm were prepared using a vibratome and maintained fully hydrated in the same medium until acquisition, without drying or fixation. Imaging was performed within 3hours of excision.

Male Wistar Albino Glaxo (WAG)/Rij rats (10 to 12 weeks, body weight ≥ 200 g, mean 250 g) were purchased from Envigo (Horst, The Netherlands), acclimatized for at least 7 d, and housed under standard conditions with food and water *ad libitum* in a temperature- and humidity-controlled room on a 12 h light/dark cycle. Animal experiments were approved by the Animal Ethics Committee of the Faculty of Medicine of Ghent University (ECD 22-41) and performed according to Belgian and European animal welfare regulations.

### Fluorescence Lifetime Imaging

Fluorescence lifetime imaging microscopy was performed using an inverted Leica Stellaris 8 FALCON confocal microscope with a DMI8-CS stand, a pulsed tunable white-light laser, and hybrid detectors operated in photon-counting mode under LAS X version 4.8.2.29567 (Leica Microsystems, Germany). The system was equipped with APO 10×/0.40 air, HC PL APO CS2 40×/1.25 glycerol, and HC PL APO 63×/1.40 oil-immersion objectives. The white-light laser was operated at a repetition rate of 80 MHz. Images were acquired at 512 × 512 pixels and 16-bit depth using a galvanometric scanner operated at 400 Hz in unidirectional mode, with a pixel dwell time of 2.8 µs, three-frame averaging, two-line accumulation, and a pinhole diameter of 1.0 Airy unit.

Fluorescence channels were acquired sequentially to minimize spectral crosstalk. Tissue autofluorescence and DOX fluorescence were acquired in a common channel, excited at 475 nm and detected from 491 to 648 nm for lifetime imaging, with the narrower 491–560 nm interval used where indicated for intensity analysis. DiBuO-aza-BODIPY fluorescence was excited at 698 nm and detected from 703 to 750 nm. Fixed tumor sections were mounted using ready-to-use VECTASHIELD Antifade Mounting Medium with DAPI (H-1200-10; Vector Laboratories, USA). DAPI fluorescence was acquired sequentially following excitation at 405 nm. Acquisition settings were held constant across conditions compared within each experiment.

### Ex Vivo NP Incubation and Matrix-Dependent Lifetime

Fresh human peritoneal sections were incubated with aza-BP NPs diluted in normal saline to 146 µg mL□¹ for 30 min at room temperature. Sections were washed twice with normal saline and imaged in imaging medium at 37 °C without fixation. Untreated tissue sections acquired during the same imaging session served as autofluorescence controls. This concentration was used to establish the tissue-associated NP reference and was not used for the subsequent penetration experiments.

We evaluated the matrix dependence of the aza-BODIPY lifetime by imaging aza-BODIPY NPs at 0.5 mg mL□¹ in deionized water, imaging medium, 25% fetal bovine serum (FBS), fresh human peritoneum, and rat tumor tissue. All measurements were acquired at 37 °C using identical settings, with five independent measurements per matrix.

### Tissue Processing and Ex Vivo Human Peritoneum Experiments

Tissue used for depth-resolved analysis was embedded in optimal cutting temperature compound (Sakura Finetek, Japan), snap-frozen, and cryosectioned at 30 µm perpendicular to the exposed surface. Sections were fixed in precooled methanol for 10 min at −20 °C and mounted using the mounting medium Vectashield (Vector Laboratories, USA).

The NP working concentration was evaluated in the previously described ex vivo EMDA model.^21^ Dye-only RL-100 NPs were tested at 73, 29.2, and 14.6 µg mL□¹ under PD or EMDA. Twelve tissue samples from three independent experiments were analyzed. Based on signal detectability across the analyzed depth, 14.6 µg mL□¹ was selected for the subsequent animal experiments.

### NP Delivery in Healthy Rats

Three groups of three rats received an IP instillation of 50 mL normal saline under sevoflurane general anesthesia. The procedural-control group received current without NPs. The PD group received dye-only RL-100 NPs at 14.6 µg mL□¹, corresponding to 0.73 mg per animal or approximately 2.9 mg kg□¹, without current. The EMDA group received the same NP dispersion with concurrent electrical treatment for 30 min. Dye-only NPs were used to assess carrier distribution without interference from DOX fluorescence.

EMDA was delivered using a customized platinum-electrode fluid-column system connected to a PHYSIONIZER 30 N2 generator (Physion s.r.l., Italy), following the previously validated configuration.^21^ Current was increased incrementally from 0 mA to the maximum tolerated 1.6 mA and held for 30 min, with continuous recording of current, voltage, and impedance. Heart rate, respiratory rate, and oxygen saturation were monitored throughout the procedure. The 50 mL instillation volume was required to establish the conductive fluid column used for electromotive transport. The procedure was performed under terminal general anesthesia, with euthanasia immediately after treatment, as its tolerability was established previously.^21^ Right and left parietal peritoneal tissues were collected and processed as described above.

### Monitoring of Intracellular DOX Release in CC531s.2 Cells

CC531s.2 rat colorectal carcinoma cells were seeded at 20 000 cells per well in 250 µL RPMI 1640 supplemented with 10% fetal bovine serum (Thermo Fisher Scientific, USA) in eight-well chambered µ-slides (ibidi, Germany). Cells were allowed to adhere overnight at 37 °C in 5% CO□. The seeding density produced approximately 30% confluence at the first imaging time point. Cells were exposed to DOX azaBP NPs or free DOX at DOX-equivalent concentrations of 2 or 20 × 10□□ M. Dye-only aza-BODIPY-labeled RL-100 NPs were applied at NP concentrations matching those used for the co-loaded formulations. A vehicle control received the corresponding volume of glycerol-containing NP storage medium. Cell cultures were observed over 24 h and imaged after 1, 5, and 24 h. Before acquisition, culture medium was replaced with imaging medium. Three independent experiments were performed, with five fields acquired per well and at least ten cells analyzed per field of view.

### Tumor-Bearing Rat Model

PM was induced by IP injection of 2.0 × 10□CC531s.2 cells in sterile phosphate-buffered saline on day 0, using the previously established syngeneic model.^21^ Body weight and general condition were assessed from day 0 to 15, and treatment was administered on day 15. Body-weight measurements are shown in Figure S6a.

Five groups of three rats were studied: DOX azaBP NPs under PD with collection at 6 h; the same formulation under EMDA with collection at 6 or 24 h; and free DOX under PD or EMDA with collection at 6 h. All treatments were administered intraperitoneally in 50 mL normal saline at a DOX-equivalent dose of 3 mg kg□¹. Meloxicam (2 mg kg□¹, subcutaneous) was administered before the procedure. Anesthesia and EMDA were performed as described for the healthy-rat experiments. At the assigned endpoint, animals were euthanized, and tumor nodules and peritoneal tissues were harvested systematically, with particular attention to the tumor-bearing omentum. Tissues were processed as described above, and parallel sections were stained with hematoxylin and eosin for confirmation of tumor identity and histology.

### Whole-Animal and Ex Vivo Near-Infrared Imaging

Carrier distribution was evaluated using an IVIS Lumina LT system (PerkinElmer, USA) under isoflurane anesthesia. Only animals receiving DOX azaBP NPs were included because free DOX emission lies outside the Cy5.5 detection window. A saline-treated animal was used to establish background fluorescence.

Animals assigned to the 6 h endpoint were imaged 1 and 6 h after PD or EMDA, whereas the 24 h EMDA group was imaged at 1, 6, and 24 h. Final imaging was performed immediately before euthanasia in lateral and supine positions using 640 nm excitation and a Cy5.5 emission filter spanning 695–770 nm. Epi-illumination, binning of 4, a 12.5 cm field of view, f/2, and an exposure time of 0.5 s were used throughout. After euthanasia, tumor nodules, tumor-bearing omentum, right and left parietal peritoneum, mesentery, liver, kidneys, spleen, and stomach were excised and imaged individually using the same settings. Average radiance was quantified from standardized regions of interest using Living Image version 4.7.4. Measurements were background-subtracted, and five scans were averaged per animal.

### Image and FLIM-Phasor Analysis

Fluorescence lifetime data were analyzed using LAS X version 4.8.2.29567 and GSLab, a MATLAB-based open-source platform for fluorescence lifetime and spectral phasor analysis.^65^ Phasor transformations were calculated at the first harmonic of the 80 MHz laser repetition frequency.^49,66^ No external lifetime calibration was applied because the Leica FALCON system performs internal calibration using the reflected excitation signal. A median-filter setting of 15 and an intensity-threshold setting of 10, as specified in LAS X, were applied identically to all phasor data.

Circular phasor masks were placed around the phasor populations of interest. Within each experiment, the center and radius of each phasor mask were fixed across the treatment groups and time points being compared. Apparent lifetimes are reported as phase lifetimes (τφ) derived from the phasor coordinates.^49,66^ For human and healthy-rat peritoneum, a circular phasor mask was first applied to extract NP-associated fluorescence, after which mean near-infrared intensity was quantified within predefined spatial regions of interest. For tumor DOX penetration, phasor plots were used to characterize the DOX-associated population, whereas the depth profiles were calculated from intensity images using a common threshold applied identically across treatment groups and depths. No lifetime-based selection was applied to the tumor penetration profiles.

Tissue blocks were oriented so that cryosectioning produced cross-sections perpendicular to the exposed surface. Penetration depth therefore represents the in-plane distance from the tissue surface and is orthogonal to the 30 µm section thickness; it does not represent axial optical imaging depth. For peritoneal tissue, mean fluorescence intensity was measured in three consecutive 100 µm regions: ROI 1, 0–100 µm; ROI 2, 101–200 µm; and ROI 3, 201– 300 µm from the exposed mesothelial surface. Triplicate sets of 300 × 100 µm² rectangles were placed perpendicular to the surface at fixed 500 µm intervals across each section. For tumor tissue, consecutive 100 µm bands were extended from the nodule surface across the full available tissue depth.

For the cell-release experiments, DOX-rich nuclear regions were segmented from the DOX channel using Cellpose.^67^ The resulting masks were applied to the DOX intensity and lifetime images to obtain the mean nuclear fluorescence intensity and phase lifetime for each nucleus. At least ten nuclei were analyzed per field. Because the analysis was restricted to nuclear regions, the measured lifetime describes the state of nuclear DOX rather than the amount of DOX remaining associated with NPs in the cytoplasm.

Tumor sections were imaged using automated tile scanning and stitching in LAS X. Each reconstructed mosaic comprised approximately 4 × 4 adjacent fields, with the number of tiles adjusted to cover the available tumor area from the nodule surface toward the core. Unlike the phasor-mask-based selection used for peritoneal NP analysis, tumor depth analysis was performed using a custom Python script applied to exported ROI data containing spatial coordinates, mean fluorescence intensity, and mean fluorescence lifetime. The algorithm identified the tissue boundary and divided the sample into successive 100 µm depth layers. ROIs were assigned to layers according to their spatial coordinates, and their mean intensity and lifetime values were averaged within each layer to generate depth-resolved profiles. Tumor DOX intensity profiles were calculated without lifetime-based ROI selection. Three animals were included per treatment group, with three tissue samples collected per animal. Replicate measurements were averaged within each tissue sample before group-level analysis. The analysis notebooks are available through the repository listed in the Data Availability Statement.

### Statistical Analysis

Data are presented as mean ± standard deviation, with normality assessed by quantile-quantile plots and the Shapiro-Wilk test. Depth-resolved intensity in ex vivo human and healthy-rat peritoneum was compared between PD and EMDA using two-way repeated-measures analysis of variance followed by Tukey-adjusted comparisons.

Tumor penetration profiles were analyzed separately for NP-delivered DOX and free DOX using generalized estimating equations with a first-order autoregressive working correlation. Depth-resolved measurements from the same imaged region were treated as repeated measures within a cluster. Depth was entered as a second-order polynomial, with treatment group, depth, and the group-by-depth interaction included in each model. Lifetime measurements were analyzed descriptively. All tests were two-sided, and p < 0.05 was considered statistically significant. Analyses were performed using R version 4.6.1 (R Foundation for Statistical Computing, Austria) and GraphPad Prism version 8.3.0 (GraphPad Software, USA).

## Supporting Information

Supporting Information is available from the Wiley Online Library or from the author.

## Use of AI tools

During the preparation of this work, the authors used ChatGPT (OpenAI) and Claude (Anthropic) to improve the writing, assist in drafting and revising selected text, and generate short Python scripts for plotting and for organizing the depth-resolved intensity data. The tools were not used to generate, analyze, or alter research data. The authors reviewed and validated all AI-assisted text and code and take full responsibility for the content of the publication.

## Data Availability Statement

The Python notebooks used for depth-resolved FLIM analysis and figure generation are publicly available at the GitHub link.

