## Supplementary Data File for "Electrically Enhanced Intraperitoneal Drug Delivery Resolved by Fluorescence Lifetime Imaging Microscopy"

### Nanoparticle Characterization

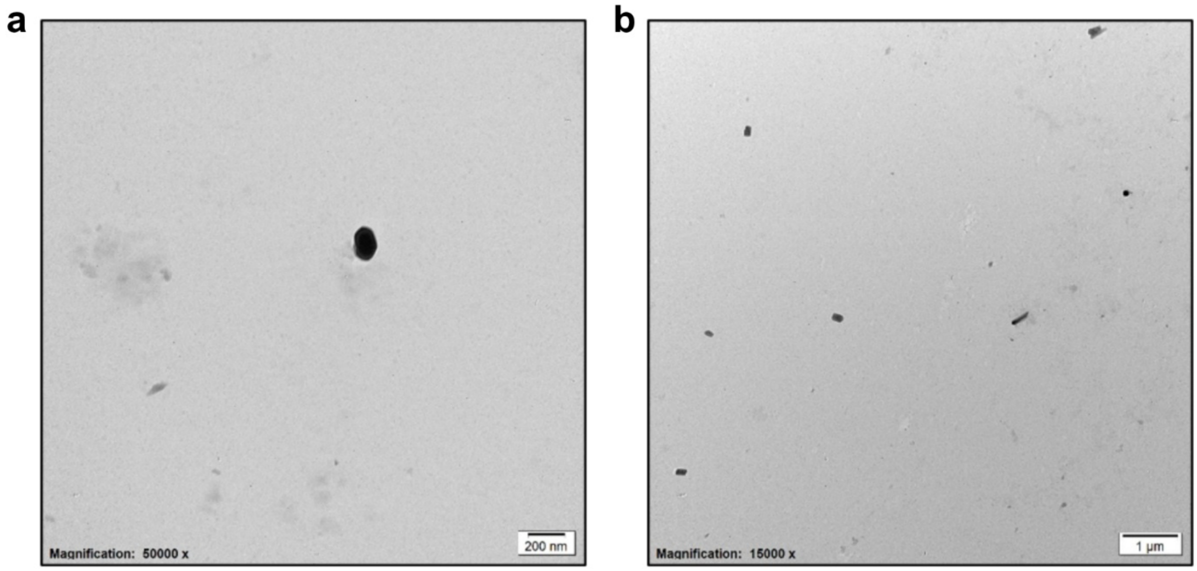


**Figure S1.** Transmission electron microscopy of aza-BODIPY-labeled RL-100 NPs without DOX. a,b) Representative TEM images from the batch used in this study. Analysis of 150 NPs yielded a mean diameter of 58.4 ± 3.5 nm (mean ± SEM). Scale bars: a) 200 nm; b) 1 µm.

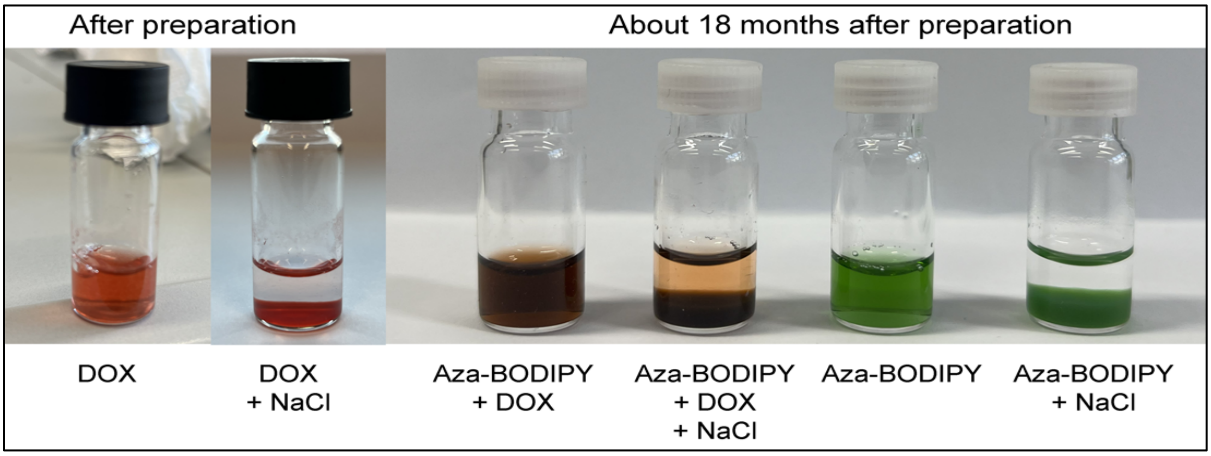


**Figure S2.** Qualitative assessment of payload retention by salt-induced precipitation. Photographs of RL-100 NP dispersions before and after addition of saturated aqueous NaCl at a volume ratio of 1:5 relative to the NP dispersion. From left to right: freshly prepared DOX-containing dispersion without and with NaCl; DOX/diBuO-aza-BODIPY co-loaded dispersion approximately 18 months after preparation, without and with NaCl; and diBuO-aza-BODIPY-only dispersion approximately 18 months after preparation, without and with NaCl.

### In Vivo Delivery in Healthy Rats

**Table S1.** Electrical parameters and physiological monitoring during intraperitoneal delivery in healthy rats. Values are means of three animals per group.

| Parameter | Control (EMDA, no NPs) | EMDA + azaBP NPs | PD + azaBP NPs |
|---|---|---|---|
| Current amplitude | 1.0 to 1.6 mA | 1.0 to 1.6 mA | Not applicable |
| Maximum voltage | 6.6 V | 6.3 V | Not applicable |
| Heart rate [bpm] | 262 | 264 | 240 |
| Respiratory rate [$min^{-1}$] | 46 | 49 | 35 |
| Oxygen saturation [%] | 99.5 | 99.1 | 99.7 |

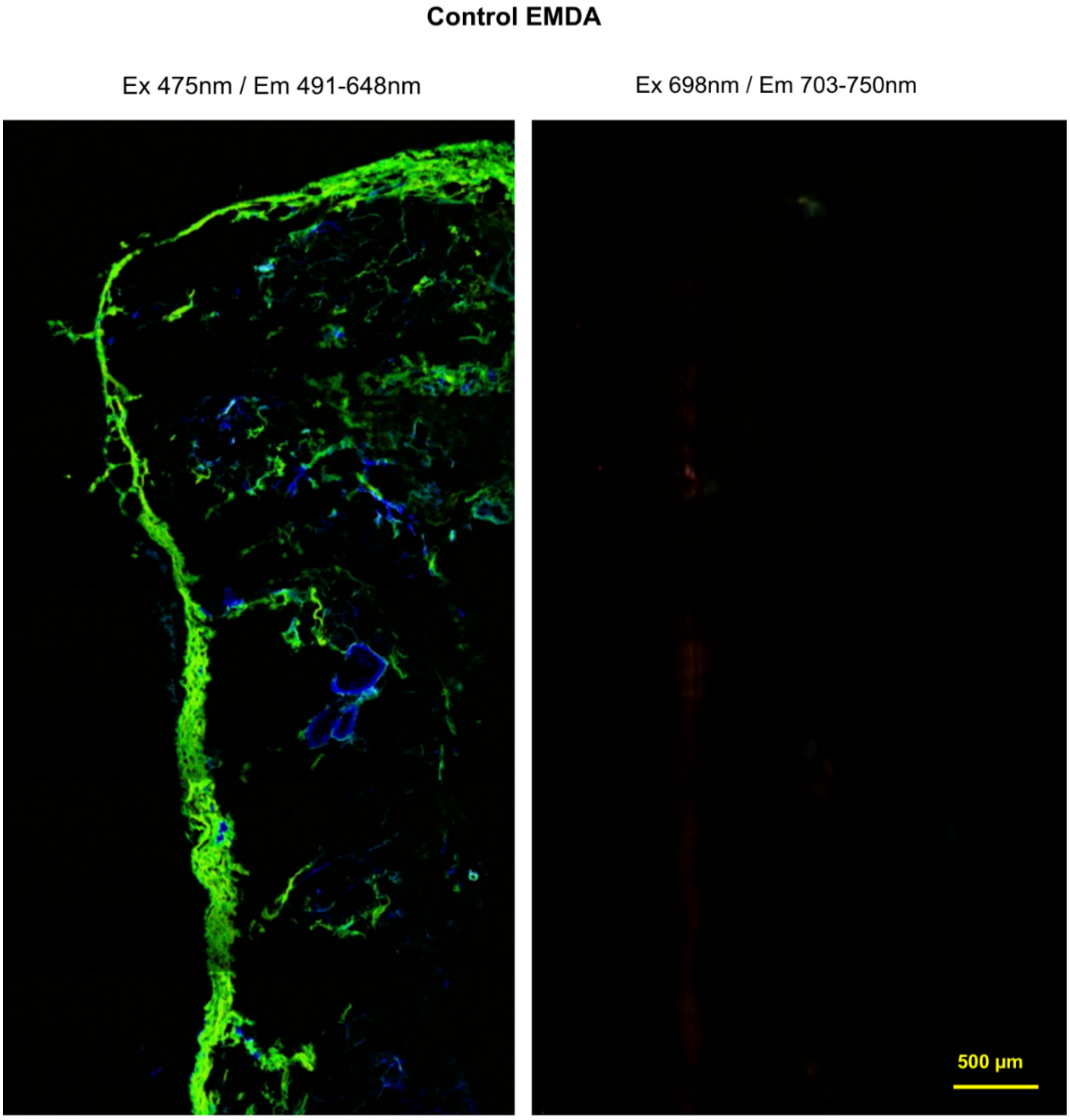


**Figure S3.** Procedural control in healthy rat parietal peritoneum. Representative tissue section following intraperitoneal saline administration with EMDA, without NPs. Images were acquired under the conditions used for NP-treated tissue in the visible (λexc = 475 nm; λem = 491–648 nm) and NIR (λexc = 698 nm; λem = 703–750 nm) channels. No discernible NIR signal was observed above background. Scale bar: 500 µm. The NIR channel is displayed within the same intensity range as the corresponding NP-treated images in the main text.

### Intracellular DOX Release

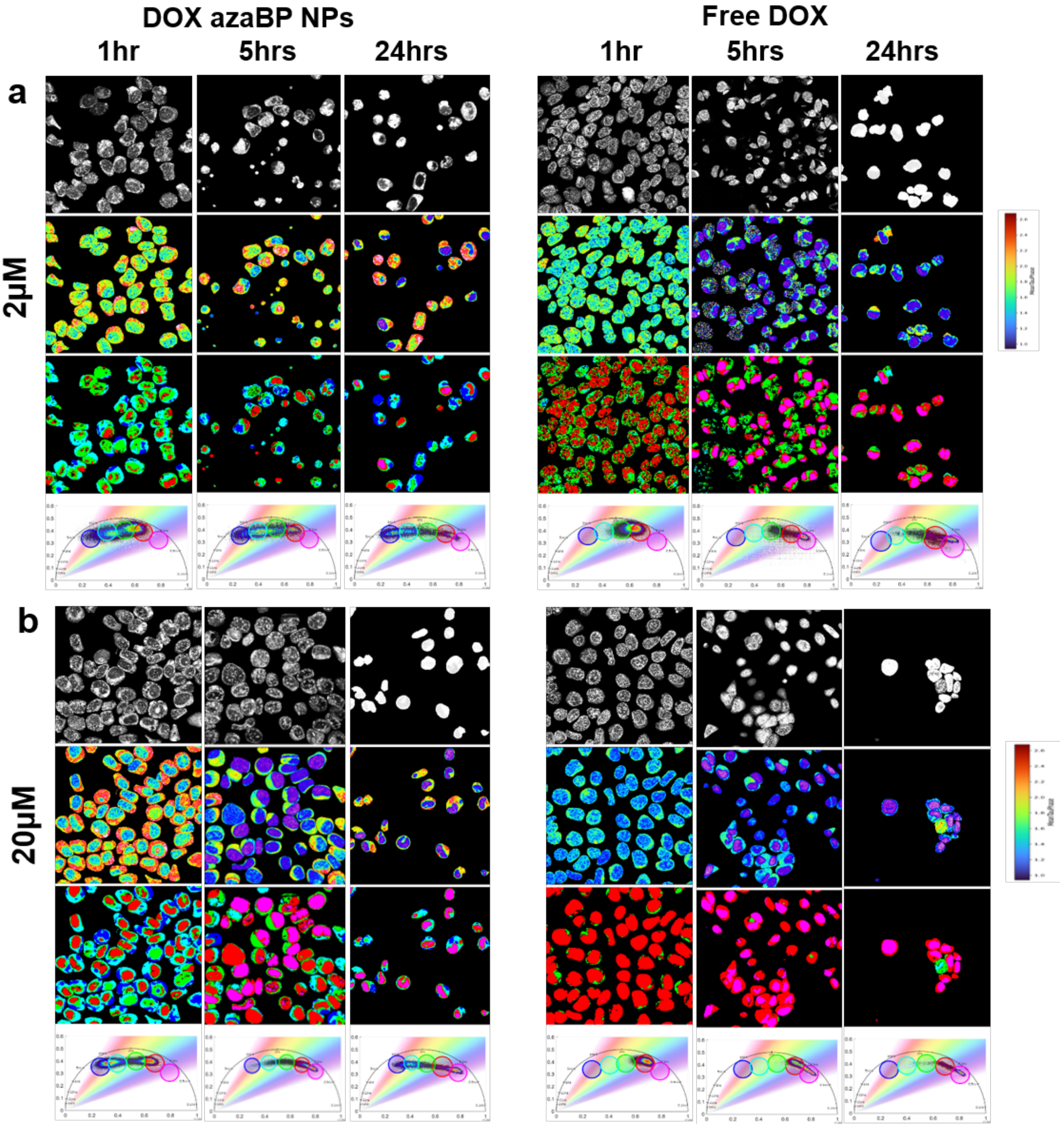


**Figure S4.** Intracellular DOX distribution and fluorescence lifetime in CC531s.2 cells. Complete image set following exposure to DOX-azaBP-NPs or free DOX at DOX-equivalent concentrations of a) 2 µM and b) 20 µM for 1, 5, and 24 h. For each condition, rows show DOX fluorescence intensity in grayscale, phase-lifetime-coded fluorescence using a common 0.95–2.70 ns scale, pixels pseudocolored according to the circular phasor masks, and the corresponding phasor plots. Scale bar: 20 µm.

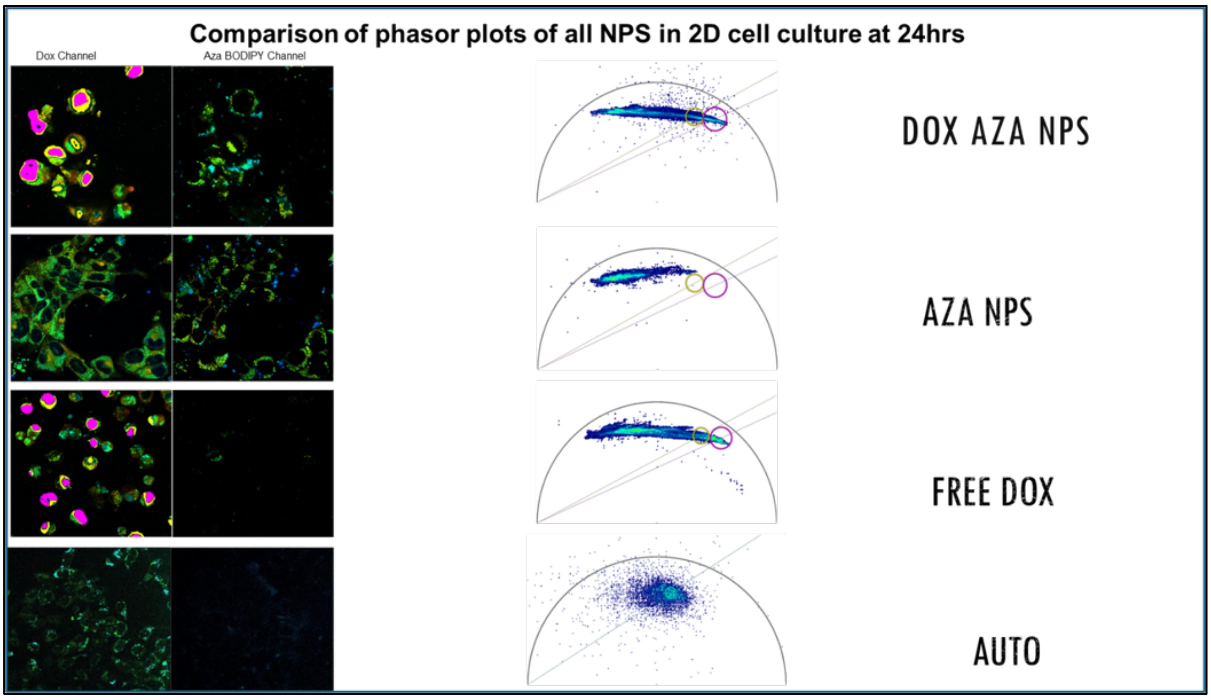


**Figure S5.** Carrier–payload separation and fluorescence-channel controls in CC531s.2 cells at 24 h, using 20 µM DOX-equivalent concentrations for the DOX-containing treatments. Representative DOX-channel and aza-BODIPY-channel images and corresponding phasor plots following exposure to DOX-azaBP-NPs, azaBP-NPs without DOX, or free DOX, together with untreated-cell controls. Carrier-associated aza-BODIPY fluorescence remained predominantly cytoplasmic, whereas DOX accumulated in nuclei. Dye-only NPs and untreated cells showed no discernible signal above background in the DOX channel under the same acquisition conditions. Images within each channel are presented within a similar intensity range for comparison.

### Tumor Study Monitoring

a

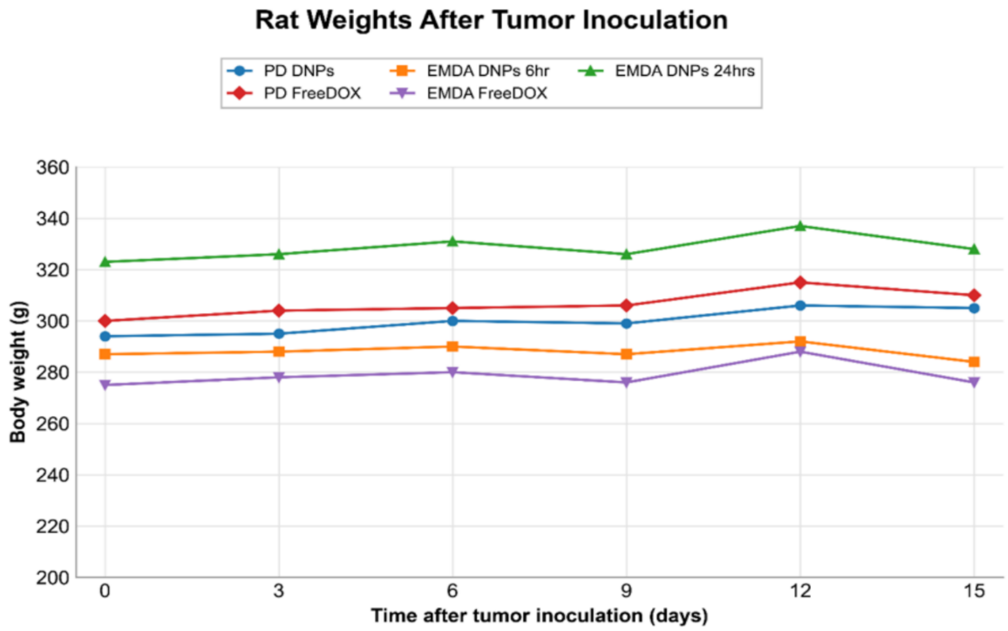


b

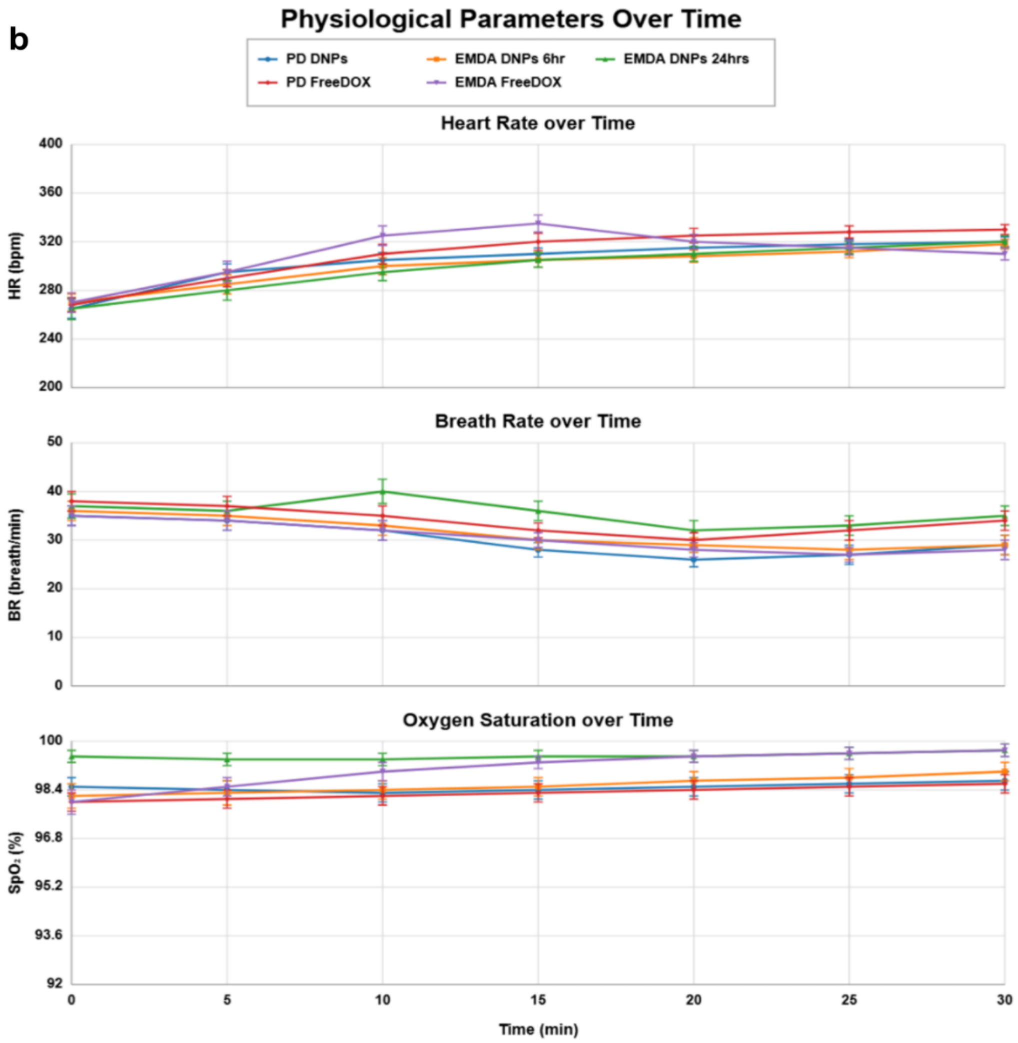

**Figure S6.** Body weight and physiological monitoring in tumor-bearing rats. a) Mean body weight following tumor inoculation, measured on days 0, 3, 6, 9, 12, and 15. b) Heart rate, respiratory rate, and oxygen saturation during the 30 min intraperitoneal treatment period. Groups received DOX-azaBP-NPs under PD or EMDA, with collection at 6 h or, for EMDA only, 24 h, or free DOX under PD or EMDA with collection at 6 h. Data are presented as mean ± SD; n = 3 animals per group.

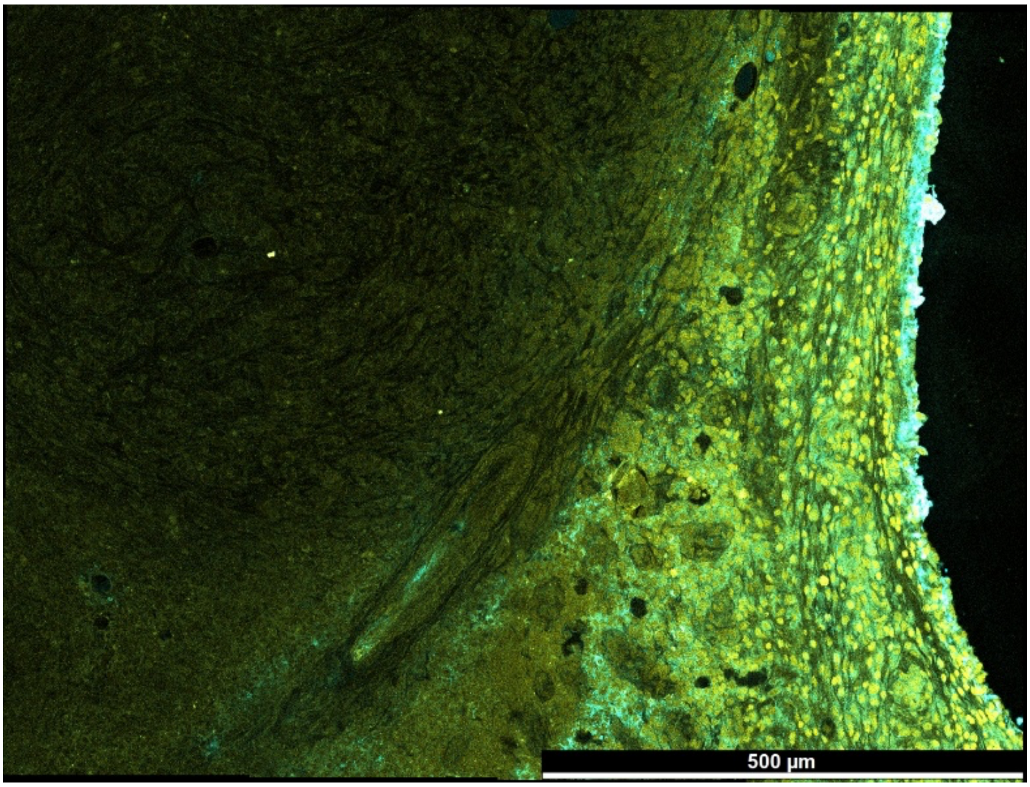


**Figure S7.** Representative mosaic image of a colorectal peritoneal tumor nodule 6 h after intraperitoneal delivery of DOX/aza-BODIPY co-loaded RL-100 NPs by EMDA. The image illustrates the spatial distribution of DOX-channel fluorescence within the tumor section and complements Figure 7 of the main text. DOX fluorescence was excited at 475 nm and detected at 491–648 nm. Scale bar: 500 µm.

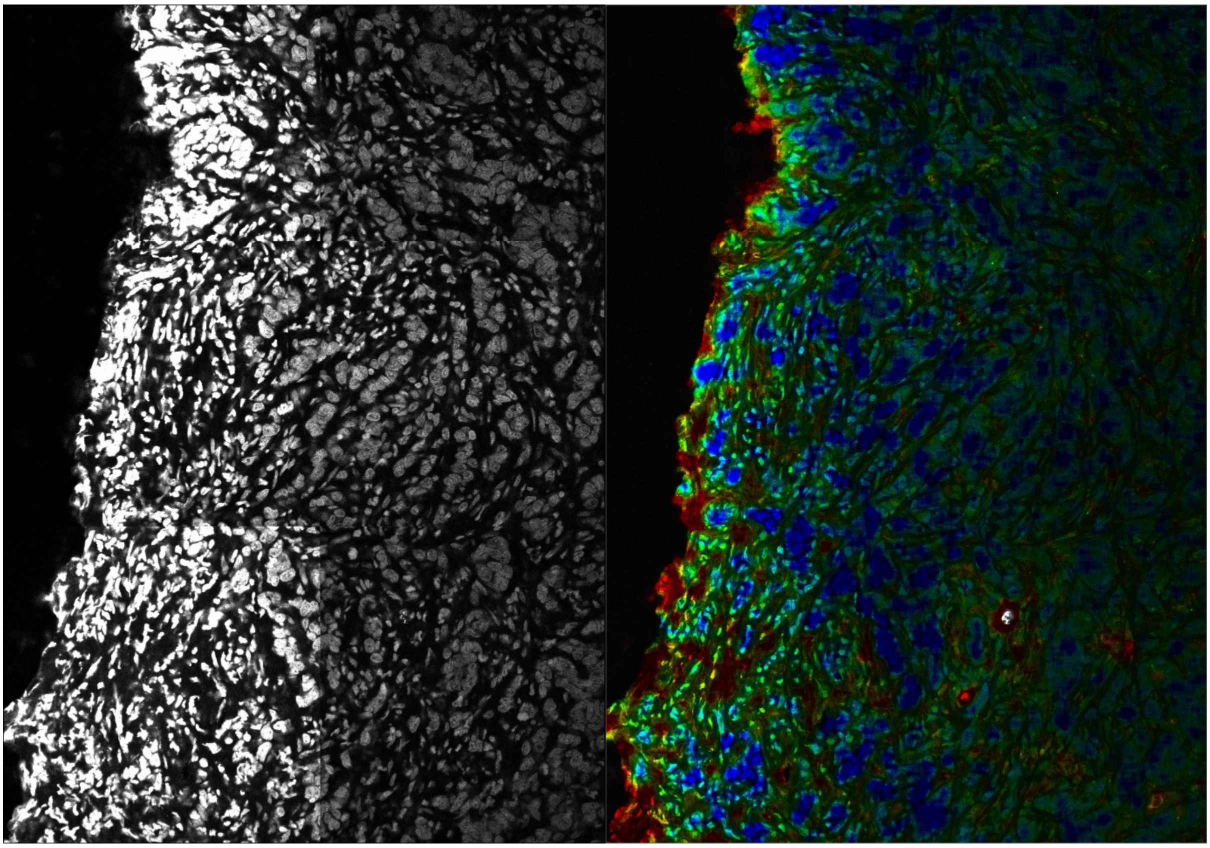

**Figure S8.** Higher-magnification view of DOX distribution at the tumor surface. DOX-channel fluorescence intensity (left, grayscale) and phase-lifetime-coded DOX fluorescence (right; λexc = 475 nm; λem = 491–648 nm) in a peritoneal tumor nodule 6 h after intraperitoneal delivery of DOX/aza-BODIPY co-loaded RL-100 NPs by EMDA.
